# A Dynamic Model of Calcium Frequency Modulation by Mitochondrial Superoxide in Non-excitable Cells

**DOI:** 10.1101/2025.07.28.667258

**Authors:** Ali Same-Majandeh, Seyed Peyman Shariatpanahi, Shahrzad Hadichgeni, Amirali Zandieh, Bahram Goliaei

## Abstract

Numerous studies have highlighted the crucial roles of cytosolic calcium transients mediated by inositol 1,4,5-trisphosphate receptor (IP3R) channels in intracellular signaling, as well as the significant modulatory effects of cytosolic redox state on these dynamics. Accordingly, developing a mathematical model that captures the core dynamics by which redox state of IP3Rs influence calcium signaling would provide a valuable foundation for simulating and predicting experimental observations. In this study, we developed a dynamic system, based on ordinary differential equations that incorporates a kinetic model for the redox dynamic of the IP3R and the role of this dynamic on the channel’s gating parameters. Using this framework, we simulated both the static and dynamic effects of mitochondria-derived superoxide fluctuations on local calcium oscillations in non-excitable cells. Numerical solutions demonstrated that the model quantitatively reproduced experimentally observed changes in local calcium oscillation frequency in response to mitochondria-derived superoxide variations. Overall, the proposed model offers a predictive tool for exploring how redox perturbations in non-excitable cells affect cellular signaling pathways through calcium frequency modulation.

## Introduction

The dynamics of free calcium in the cytosol of cells—appearing as global oscillations, propagating waves, or localized oscillations—has significant physiological importance across a wide range of cell types, including excitable cells such as neurons and cardiac cells, and non-excitable cells (many of other somatic cell types) (1–3). Under conditions of cellular activity, these dynamics dominantly appear as global oscillations or propagating waves in the cytosol, controlling vital signaling processes such as cell differentiation, gene expression and secretion of hormones or other substances (4–6). In contrast, in resting conditions, calcium dynamics in the cytosol often appear as localized fluctuations—known as calcium flashes, sparks, or puffs—and play a critical role in maintaining the stability of home-keeping functions, particularly mitochondrial function and thus energetic stability of cells (7–9). This role has been observed in both excitable and non-excitable cells (10). Meanwhile, numerous studies have demonstrated that the frequency of calcium oscillations, rather than solely their amplitude, is the key determinant of their physiological functions (11,12).

On the other hand, several studies have indicated that cellular reactive oxygen species (ROS), especially mitochondria-derived ROS, exhibit localized oscillations under pathological or even physiological conditions (13–15). For instance, research on a variety of cell types revealed that a single mitochondrion generated pulses of superoxide in the surrounding cytosol under resting cellular conditions (16).

Parallel to above studies, the relationship between ROS and calcium dynamics has been investigated for decades (17). For example, it has been shown that calcium dynamics is influenced by mitochondrial ROS dynamics or the redox state dynamics of the cytosol, and alterations in the concentration of ROS or changes in the reducing capacity of the cytosol modify calcium dynamics (18). Particularly, observations in this context have shown that an increase in cytosolic ROS, under resting conditions that concentration of inositol 1,4,5-trisphosphate (IP3)—which stimulates calcium channels in the endoplasmic reticulum (ER) membrane known as inositol 1,4,5-trisphosphate receptors (IP3R)—is at basal levels, can initiate abnormal global calcium oscillations or increase the frequency of calcium flashes (19–22). Interestingly, a recent study on local mitochondrial calcium dynamics demonstrated triggering mitochondrial superoxide production modulated the frequency of calcium oscillations, and influenced mitochondrial quality control and mitophagy in cellular models of Parkinson’s disease (23).

Indeed, studies of the molecular structure of IP3Rs have revealed that these channels, as key components of the calcium dynamics toolbox, have numerous thiol groups (in the structure of cysteine residues) on their cytosolic side and thus can be sensitive to redox changes (24–29). In fact, oxidation of these groups by a wide range of oxidants, increases the open probability of IP3Rs at a constant IP3 concentration or the duration of open state of the channels in each pulse (30,31).

Considering the aforementioned studies, which highlight the importance of calcium dynamics, various mathematical models based on systems of differential equations (ODE/PDE), or agent-based models have been developed to deepen our understanding of this dynamics and simulate both global calcium oscillations or local calcium flashes (32–46). Among these, some have specifically aimed to simulate the effect of mitochondrial ROS dynamics on the frequency of calcium flashes (45,46). However, dynamical models based on compartmentalization of different cytoplasmic regions face two fundamental issues: First, the basis for compartmentalization and the determination of species diffusion rates between non-membrane-bound compartments, such as the bulk cytosol and the microdomains of cytosol near the mitochondria, are not strictly based on geometric and physical principles but are phenomenologically determined to reproduce experimental observations. Consequently, a physical basis for determining their quantitative values is lacking. The second and more important issue is that in models developed to simulate the effect of ROS on calcium dynamics, the effect of ROS is mainly applied directly to the open probability of IP3R channels (45,46). Whereas, the open probability of IP3R channels, according to existing models, is itself a variable dependent on several parameters and has a complex dynamic (47). Therefore, the rational role of IP3R channel redox dynamics on the parameters of the open probability of IP3Rs is also missing.

Therefore, in the present study, we pursue two main objectives:

First, to adapt existing models for calcium and mitochondrial superoxide dynamics with the geometry obtained from microscopic images of these dynamics and images of the microdomains of cytosol near single mitochondria (especially the cytosolic region between a single mitochondrion and ER mitochondria-associated membranes abbreviated as MAMs) (48), and by considering a physical basis for the quantitative values of diffusion rates between different compartments.

Second, and more importantly, to develop a kinetic model for the redox dynamics of IP3R channels and a model for the dependence of IP3R channel open probability and dynamics on the redox state of these channels.

Achieving these two goals enables us to construct a more realistic computational model of the effects of ROS on calcium dynamics under both physiological and pathological conditions in non-excitable cells, and ultimately to predict how changes in key chemical or geometrical parameters may influence calcium signaling.

## Methods

### Compartmentalization

Our modeling was based on a system of ordinary differential equations (ODEs). The model was developed with six compartments: 1) the matrix of a single mitochondrion (V_M_), 2) the cytosolic region between a mitochondrion and adjacent ER membrane known as MAM (V_MAM_), which we refer to briefly as the MAM cytosol, 3) the peripheral cytosolic microdomain of a single mitochondrion (V_N_) excluding V_MAM_, which hereafter we refer to briefly as the near cytosol, 4) the bulk cytosol (V_B_), 5) the lumen of endoplasmic reticulum adjacent to a mitochondrion (V_EN_), and 6) the lumen of endoplasmic reticulum within the bulk cytoplasm (V_EB_) (Fig 1b).

**Fig 1:**
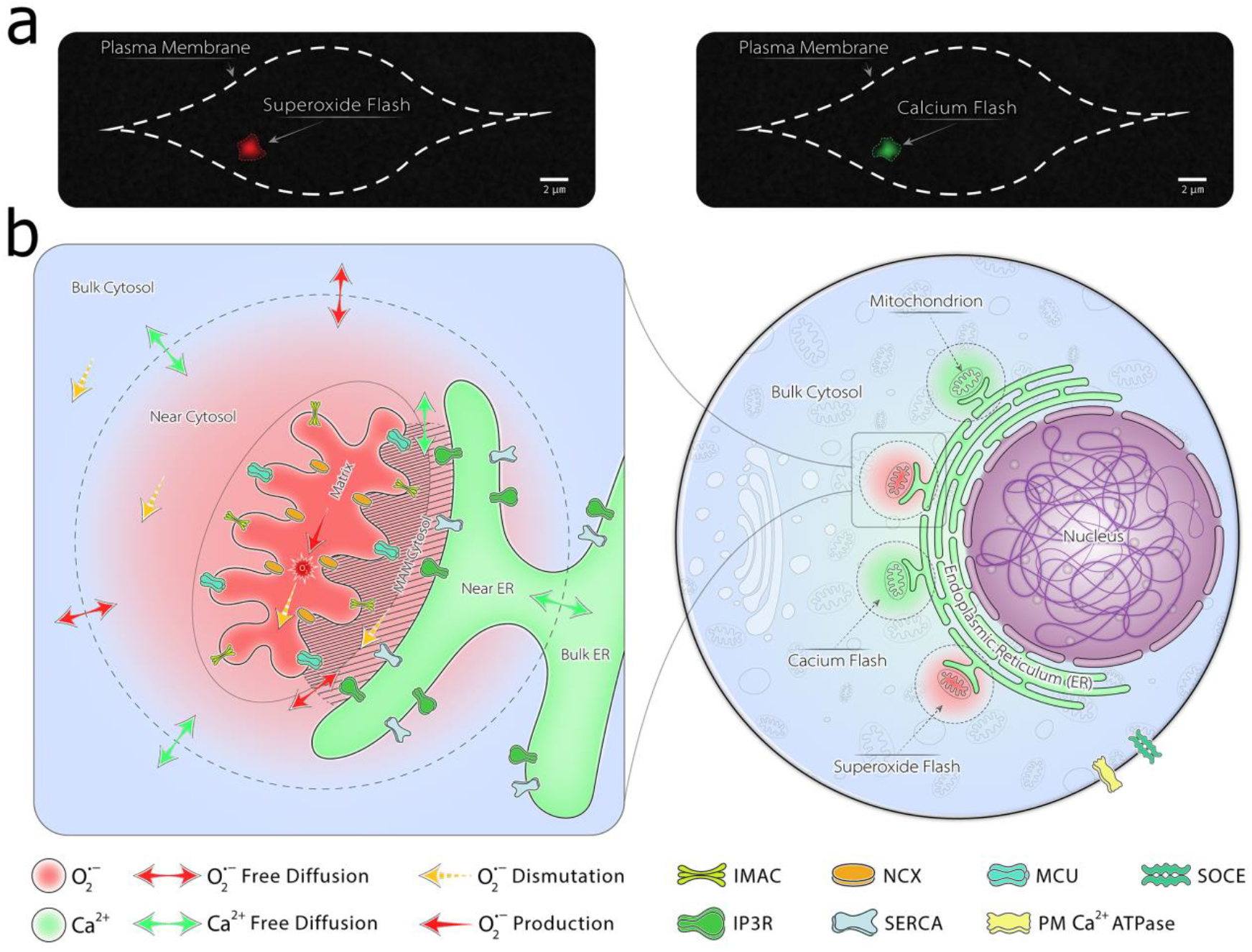
The compartments considered in the model. (a) A conceptual illustration mimicking confocal fluorescence microscopy of superoxide (left) and calcium (right) flashes. (Note: These images are stylized renderings intended solely for conceptual visualization to provide biological context, and do not represent actual experimental micrographs) (b) Schematic of compartments and fluxes in the model. The full (non-abbreviated) names of the channels, pumps, and transporters are mentioned later in the Methods section

The near cytosol (V_N_) defined based on the effective diffusion distance of superoxide following each mitochondrial superoxide flash. This region is modeled as a spherical volume surrounding a single mitochondrion, with a radius equal to the effective diffusion radius of superoxide flash (R_eff,S_).

Based on fluorescence microscopy (16), the effective diffusion radius of superoxide is estimated to be approximately 1 μm. According to the random walk model of diffusion, this radius can be derived from Eq 1:

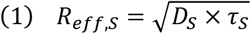

Where D_S_ is the cytosolic diffusion coefficient of superoxide and τ_S_ is superoxide half-life in the cytosol. The value of τ_S_ can be estimated based on the enzymatic degradation of superoxide using the ratio of Michaelis–Menten constant of cytosolic superoxide-dismutase enzyme to its maximum speed (Eq 2):

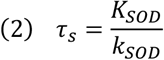

The diffusion coefficient of hydrogen peroxide in a gel-like medium is reported as 370 μm^2^/s (49). Due to the radical and ionic nature of superoxide, its diffusion coefficient is assumed to be significantly lower than that of peroxide, and is taken as 10 μm^2^/s. Therefore, to reproduce observations in fluorescence microscopy the value of τ_S_ is assumed to be 0.1 s. Finally, the volume of the surrounding cytosol estimated as the volume of a sphere with radius R_eff,S_ (Fig S2 and Eq S4).

A detailed description of the compartmentalization and geometric calculations of the model is provided in Supplementary Information.

### Dynamic Equations

#### Superoxide Dynamics

In the present study, we employed the model developed by Yang et al. (13), which incorporates ROS induced ROS release (RIRR), to describe the dynamics of superoxide in the mitochondrial matrix and cytosol. Due to the removal of the intermembrane space and the addition of the MAM cytosol compartment, we adapted the original model by Yang et al. (13) to accommodate this new compartmentalization.

The coefficients α, β, γ, δ, and ε in dynamic equations represent the volumetric ratios between different compartments (Eq S5–S9).

Here are the dynamic equations of superoxide evolution:

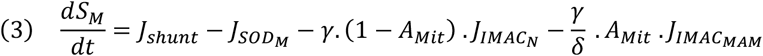

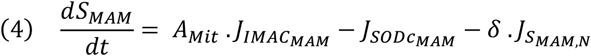

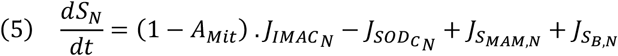

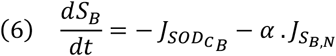

And for the mitochondrial membrane potential:

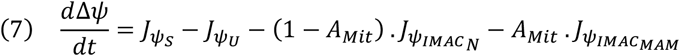

The algebraic definitions of the cytosolic superoxide dismutation rate (J_SOD,C_), mitochondrial superoxide dismutation rate (J_SOD,M_), superoxide leakage through mitochondrial inner membrane anion channels (IMAC) (J_IMAC_), the rate of mitochondrial membrane potential production by cellular respiration (J_ψ,S_), baseline membrane potential leakage rate (J_ψ,U_), and membrane potential leakage due to IMACs opening (J_ψ,IMAC_) were all adopted from the model of Yang et al. (13), and are detailed in Supplementary Information (Eq S17–S25). The nested subscripts of the fluxes—namely M, MAM, N, and B—indicate different compartments: M refers to the mitochondrial matrix, MAM denotes the MAM cytosol, N represents the near cytosol, and B stands for the bulk cytosol.

In contrast to previous models that employ phenomenological constants for the free diffusion rates of superoxide between the MAM cytosol and the near cytosol (J_S,MAM,N_), and between the near cytosol and the bulk cytosol (J_S,B,N_), we estimated these constants based on physical approximations using integrated forms of Fick’s law. The details of these estimations are provided in Supplementary Information (Eq S26–S30).

The coefficients A_Mit_ and A_ER_ (A_ER_ will be used in the calcium equations) represent the ratios of the membrane surface areas of a mitochondrion and the endoplasmic reticulum that are associated with the MAM cytosol to the total surface area of each organelle membrane (Eq 8 and 9):

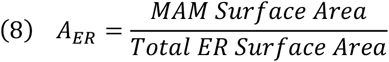

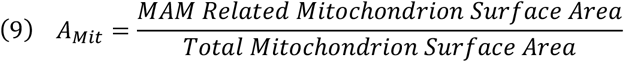

Coefficients defined as such are sufficient for determining the fluxes through channels, pumps or transporters that are uniformly distributed across the membranes of these organelles. However, for channels, transporters or pumps that are non-uniformly distributed across the membranes of the mitochondria or the ER (as will be introduced), using these ratios alone will be incorrect.

#### Calcium Dynamics

We used open-cell condition and Class 1 dynamics of calcium in our model that means calcium oscillations can occur at constant concentrations of IP3 as agonist of IP3Rs (44).

IP3R channels on the endoplasmic reticulum (ER) membrane accumulate at the MAM region through protein linkages with voltage-dependent anion channels (VDACs) located on the outer mitochondrial membrane (50). This leads to a non-uniform calcium flux into the mitochondrial matrix via mitochondrial calcium uniporters (MCUs), which reside in the inner mitochondrial membrane. As a result, the calcium flux per unit mitochondrial membrane area is elevated in the MAM region. Therefore, using only A_ER_ (for IP3Rs) and A_Mit_ (for MCUs) is insufficient to account for the non-uniform distribution of these channels across membrane regions. Thus, the additional coefficients are defined as follows:

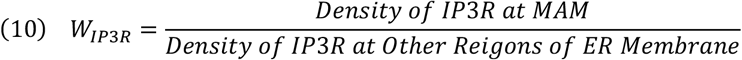

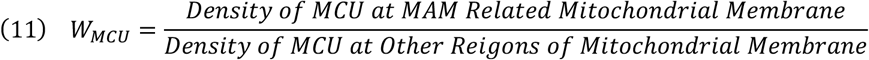

The equations governing the free calcium dynamics in the bulk cytosol (C_B_), the endoplasmic reticulum within the bulk cytoplasm (Ce_B_), the near cytosol (C_N_), the endoplasmic reticulum adjacent to a mitochondrion (Ce_N_), the cytosol within the MAM region (C_MAM_), and the mitochondrial matrix (C_M_) are as follows:

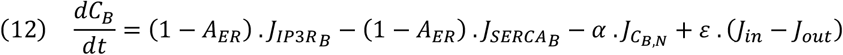

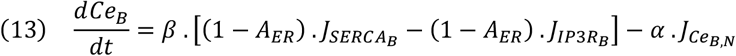

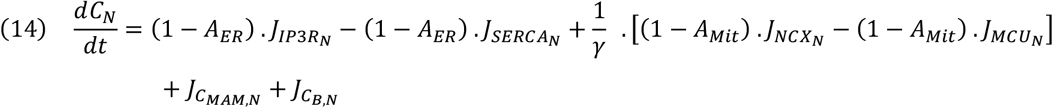

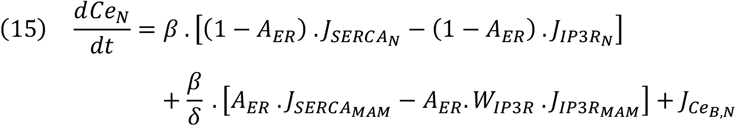

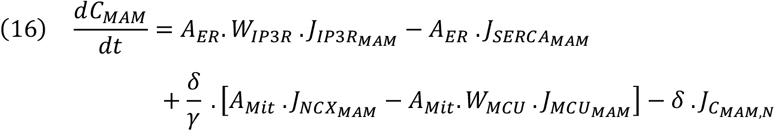

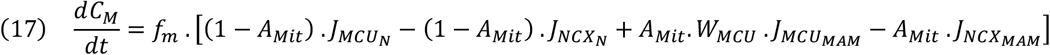

The algebraic definitions for the sarco/endoplasmic reticulum calcium ATPase (J_SERCA_), calcium influx and efflux across the plasma membrane (J_in_ and J_out_), calcium flux through IP3R channels (excluding the additional equations proposed in this study) are based on the model by Sneyd et al. (44), and are provided in Supplementary Information (Eq S49–S59). Furthermore, the equations governing calcium flux through mitochondrial calcium uniporter (J_MCU_) and through the mitochondrial sodium–calcium exchanger (J_NCX_) are based on the models by Moshkforoush et al. (38) and Wacquier et al. (36) (Eq S42-S45). The additional equations governing free diffusion of calcium between MAM cytosol and near cytosol (J_C, MAM, N_), between near cytosol and bulk cytosol (J_C,B,N_), and between ER lumen in near cytoplasm and bulk cytoplasm (J_Ce,B,N_) are also provided in Supplementary Information (Eq S46-S48).

The equations governing the time-dependent variable h for open probability of IP3R channels located at the MAM region, ER membrane adjacent to mitochondria, and at the ER membrane in bulk cytoplasm are as follows (44):

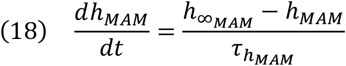

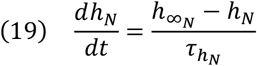

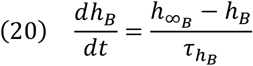

The algebraic definitions of h_∞_ and τ_h_ were adopted from Sneyd et al (44). (Eq S57 and S59).

All fluxes related to calcium and superoxide dynamics are illustrated in Fig S1.

#### IP3R redox dynamics

Experimental observations indicate that oxidation of IP3Rs increases their sensitivity to their stimulant, IP3. However, this oxidative effect on calcium dynamics through IP3R diminishes at very high IP3 concentrations (28,51,52). This strongly suggests that redox modulation of IP3R gating dynamics is primarily mediated by changing affinity of the channels for IP3. Therefore, we assumed that superoxide-induced oxidation of IP3R channels directly decreases the apparent dissociation constant of the channel for its ligand (IP3), thereby enhancing the channel’s open probability.

To model this effect, we first defined a quantity that shows oxidation extent of IP3R channels (Eq 21):

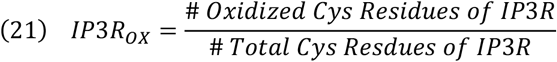

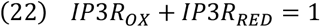

The variable IP3R_OX_ denotes the fraction of channel thiols in the oxidized state. Given the presence of multiple redox-sensitive thiols on IP3R channels (28) and assuming the cluster of IP3Rs as a single giant IP3R, we treated this fraction as a continuous variable to enable modeling its dynamics using ordinary differential equations. Due to the steady rates of IP3R channel expression and degradation throughout the simulation, the concentration of IP3R channels, and thus the denominator of Eq 21, is assumed to remain constant.

Subsequently, the K_P_, which represents the half maximal concentration of IP3 for feedback on the IP3R channels as defined in the model by Sneyd et al. (44) (Eq S58) and theoretically shows apparent dissociation constant of IP3 from IP3Rs, was expressed as a function of the oxidation fraction of the channel (IP3R_OX_) (Eq 23):

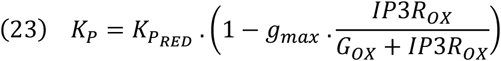

In above equation, K_P,RED_ represents the maximum value of K_P_ corresponding to the fully reduced state of the IP3R channels. The parameter g_max_ indicates the maximum decrease in K_P_ resulting from complete oxidation of the channel, while G_OX_ denotes the oxidation fraction of the channel at which K_P_ is decreased by half of g_max_ (i.e., the half-effective oxidation level to reduce K_P_). Fig S6 illustrates the relationship between K_P_ and the oxidized fraction of the channels (IP3R_OX_), as described by Eq 23.

To model the redox dynamics of IP3R channels under the oxidative effect of superoxide, we employed a simple kinetic model based on the scheme shown in Fig 2 and Eq 24-26 for the channels located at the MAM (IP3R_OX,MAM_), the ER membrane adjacent to mitochondria (IP3R_OX,N_), and the ER membrane in the bulk cytoplasm (IP3R_OX,B_).

**Fig 2:**
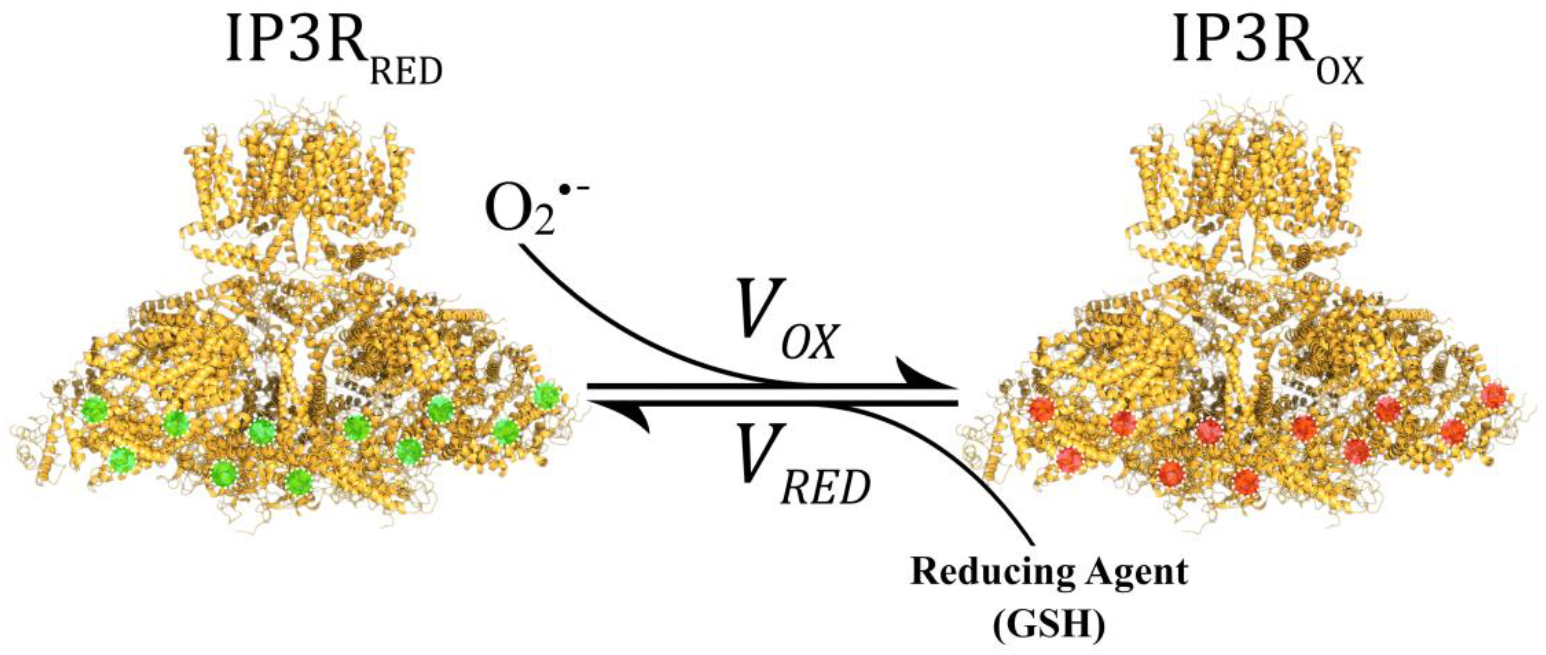
Redox reaction scheme of IP3R with superoxide and a reducing agent such as glutathione (GSH). The hypothetical reduced sites (green dots) and oxidized sites (red dots) on the channel are shown for illustrative purposes only and do not represent actual redox-active residues.

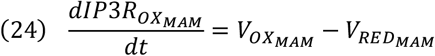

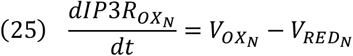

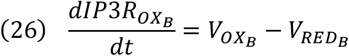

In the present model, we assumed that the reversible oxidation of the IP3R channel by superoxide follows a mass-action scheme; accordingly, the oxidation rate of the IP3R channels (V_OX_) was modeled as Eq 27. On the other hand, the reduction of the IP3R channels (V_RED_) was considered to occur enzymatically, based on a sequential reaction scheme. Furthermore, we assumed that the concentration of the cytosolic reducing agent (e.g., glutathione) remains at a level such that its potential dynamics, influenced by the ROS dynamics, do not alter the rate of enzymatic reduction of IP3R channels. Eq 27 and 28 describes the oxidation and reduction rate of the IP3R channel under these assumptions, respectively:

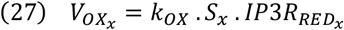

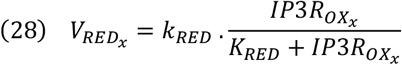

In above equations, k_OX_ is the first-order rate constant for oxidation of IP3Rs by superoxide, k_RED_ is the maximum reduction rate of oxidized IP3Rs, and K_RED_ is the fraction of oxidized IP3Rs for half-maximum speed of reduction. The subscript *x* indicates that these equations are applied to the IP3Rs in all three regions: MAM cytosol, near cytosol (N), and bulk cytosol (B).

### Model Parameters

Except where explicitly stated in the text, the remaining model parameters were either adopted from previous models or adjusted to reproduce experimental observations. The complete list of parameters used in the model is provided in Table S1.

### Numerical Solution and Sensitivity Analysis of the Model

The numerical solution of the system of ordinary differential equations was carried out using the SciPy package in Python3. Due to the inherent stiffness of the system—resulting from the oscillations in calcium and superoxide concentrations—an implicit integration method, namely the Radau algorithm, was employed to ensure numerical stability. The complete simulation script, along with the code for plotting calcium and superoxide oscillations, is publicly available on GitHub: https://github.com/Ali-Same-Majandeh/Calcium-Frequency-Modulation.

To conduct a global sensitivity analysis (GSA) of the model parameters, we employed the Regional Sensitivity Analysis (RSA) framework, based on Monte Carlo Filtering (MCF). Accordingly, 100,000 parameter sets encompassing all 52 parameters were generated via Latin Hypercube Sampling (LHS), assuming uniform prior distributions within a ±40% variation range. Following the classification of model outputs into behavioral and non-behavioral sets (discussed in Results & Discussion), the parameter sensitivities were scored using the Kolmogorov-Smirnov (KS) statistic. Both the LHS sampling procedure and the subsequent RSA/KS statistical evaluations were implemented utilizing the SALib package in Python.

## Results & Discussion

### Model Validation

The model developed in this study comprises three main components: a model for calcium dynamics, a model for superoxide dynamics, and a dynamic model describing the effect of superoxide on calcium dynamics. The first two components are primarily adopted from previous studies. Therefore, to validate behavior of these two components within the proposed compartmentalization framework, bifurcation analyses were performed for both calcium and superoxide dynamics (Fig 3 and 4). For the bifurcation analysis of calcium dynamics, the parameter k_shunt_, which represents the constant rate of superoxide production in the mitochondrial matrix, was set approximately to zero in order to examine calcium bifurcation independently of superoxide influence (Fig 3).

**Fig 3:**
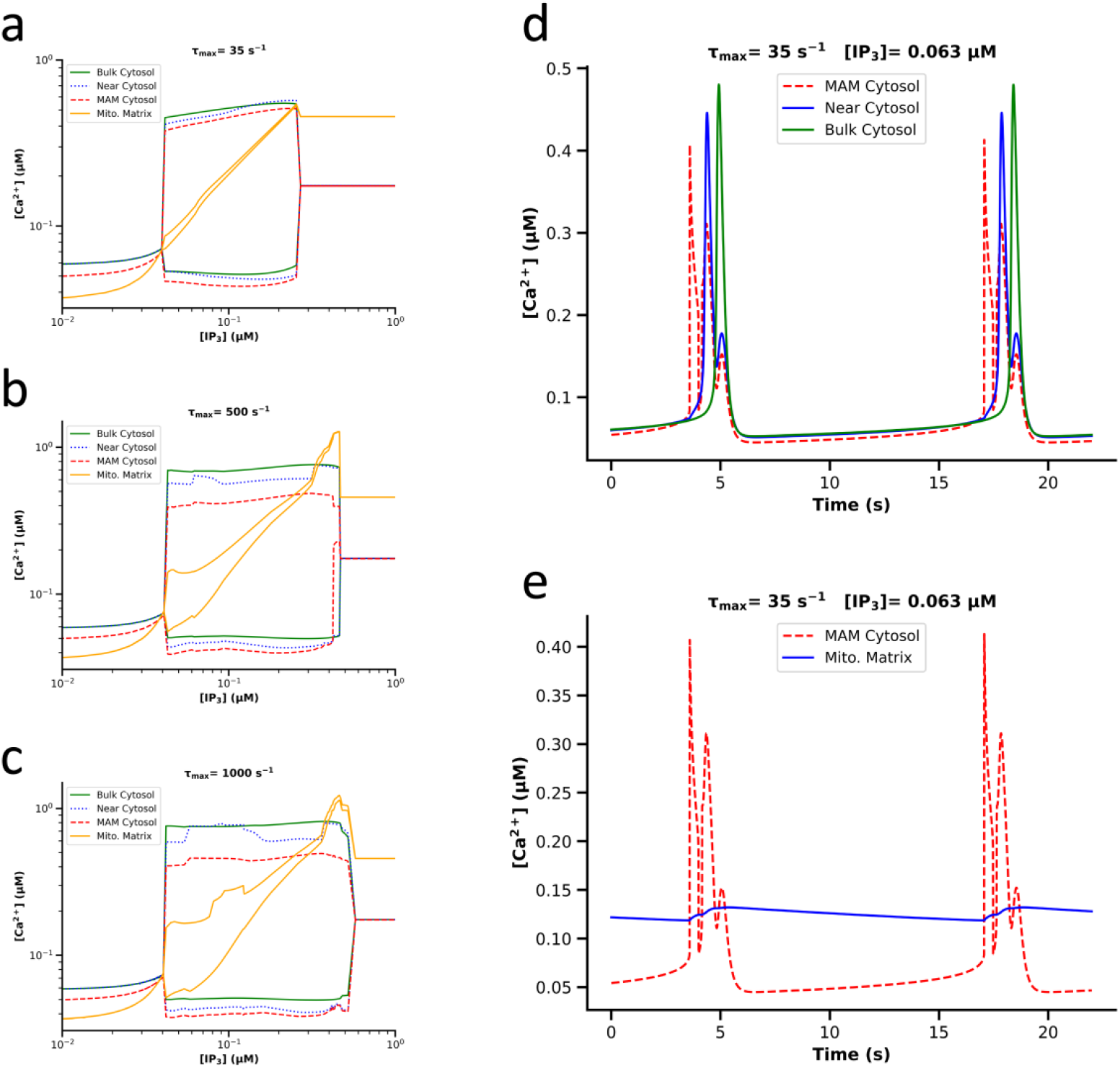
Bifurcation analysis of calcium dynamics and the calcium time-series in different cellular compartments. (a), (b), And (c) depict the bifurcation analysis of calcium based on varying concentrations of the IP3, at different values of the parameter τ_max_ (35, 500, and 1000 s^−1^, respectively). τ_max_, according to the model by Sneyd et al. (44), governs the intrinsic frequency of IP3R channel activity across different cell types. (d) And (e) show the calcium time-series in the bulk cytosol, near cytosol, MAM cytosol, and mitochondrial matrix at an IP3 concentration that simulates cellular activity conditions. A logarithmic scale was used in (a), (b), and (c) to enhance the clarity of calcium dynamics.

**Fig 4:**
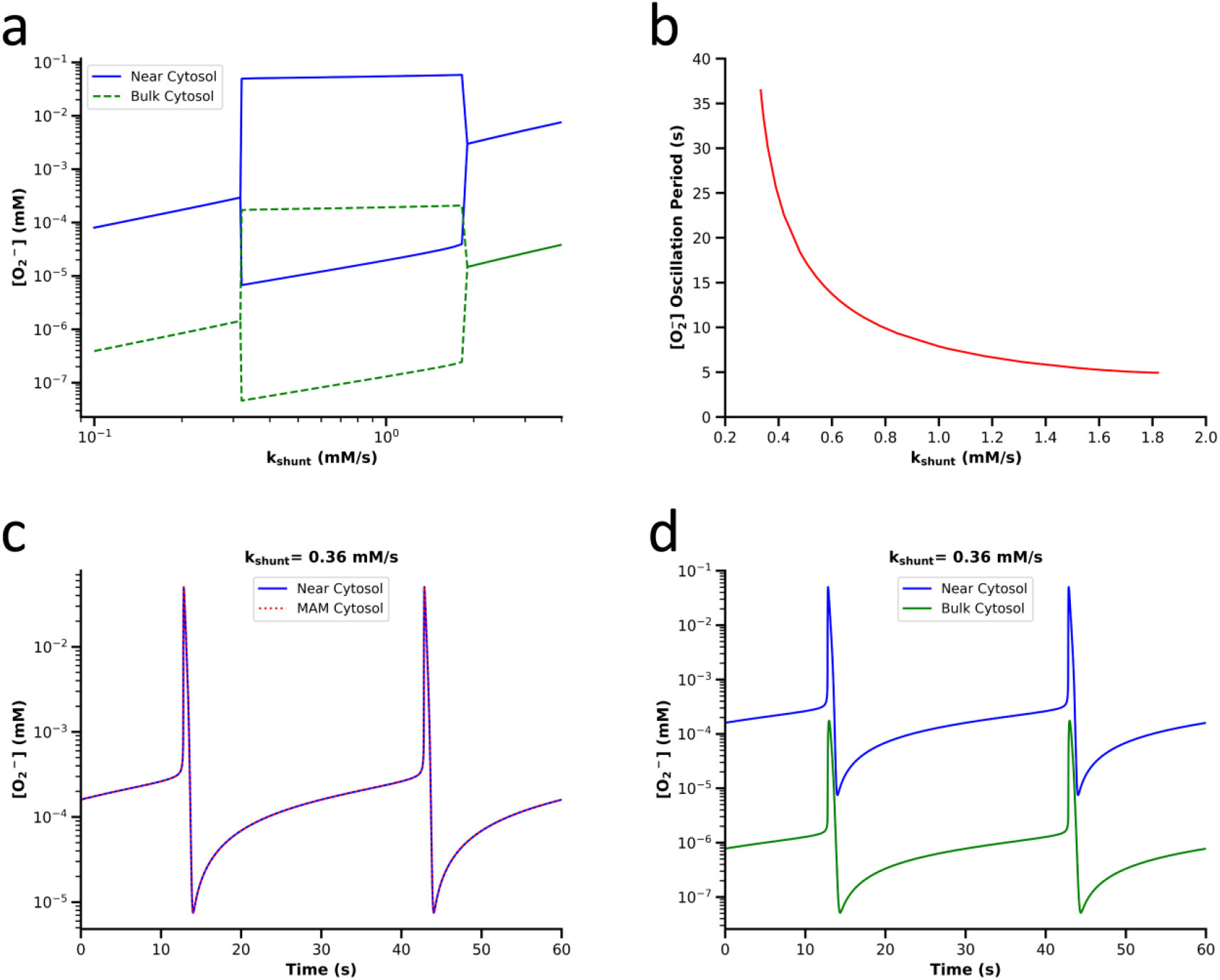
Bifurcation and oscillation frequency of superoxide dynamics. (a) Presents the bifurcation of superoxide based on the superoxide production rate in the mitochondrial matrix (k_shunt_). (b) Illustrates the frequency of superoxide oscillations as a function of the superoxide production rate within the oscillator range of k_shunt_. (c) And (d) show the time-series of superoxide in the MAM cytosol, near cytosol, and bulk cytosol at a specific k_shunt_. A logarithmic scale has been used in (a), (c), and (d) to enhance the clarity of superoxide dynamics.

The calcium bifurcation in the bulk cytosol and the near cytosol (Fig 3a-3c) is similar to the calcium bifurcation reported in the study by Sneyd et al. (44), indicating that the geometry and mitochondrial calcium fluxes implemented in the present study do not disrupt the calcium dynamics reported in previous studies or in cells with varying intrinsic calcium frequencies (by varying τ_max_). Furthermore, Fig 3a-3c demonstrate that the mitochondrial matrix calcium concentration in the current model falls within the range reported in various experimental studies and simulations (36,37,53). Examination of the calcium time-series in Fig 3d reveals that during each global calcium pulse under cellular activity conditions (i.e., when IP3 concentration in the cytosol is elevated), IP3 receptor channels (IP3Rs) in the MAM region open first, followed by calcium channels in the ER regions adjacent to the mitochondria, and finally the ER calcium channels in the bulk cytosol. The cause of this sequence and the related discussion will be addressed in the prediction section.

The markedly low range calcium dynamics in the mitochondrial matrix, shown in Fig 3e, results from the substantially higher calcium buffering capacity of the mitochondrial matrix compared to the cytosol and are consistent with previous observations and simulations (36,38,53). Additionally, the higher basal calcium concentration in the mitochondrial matrix relative to the cytosol—observable both in the calcium bifurcation analysis (Fig 3a-3c) and in calcium oscillation at each IP3 level (Fig 3e and 1d)—is consistent with experimental findings and prior modeling studies (36–38,53).

Note that the peculiarities observed in the bifurcation behavior of calcium in the near cytosol and the MAM cytosol (Fig 3b and 3c) seem due to the strong influence of perturbations originating from diffusional terms, particularly from the bulk cytosol, especially at high values of τ_max_ (See Fig S8). This is supported by the fact that such anomalous features are almost entirely absent in the calcium bifurcation within the bulk cytosol (Fig 3a-3c).

Fig 4a and 4b demonstrate that the superoxide model used in our study—an adaptation of Yang et al.’s model (13) based on the different geometry and compartmentalization of the present model—successfully simulates the amplitude and frequency of superoxide oscillations reported in Yang et al.’s model (13). Also, the present model captures the several-orders-of-magnitude difference in superoxide concentration between the mitochondria adjacent cytosol (MAM cytosol plus near cytosol) and the bulk cytosol, as expected (Fig 4a and 4d). Additionally, due to the relatively high diffusion rate of superoxide compared to the volume of the MAM cytosol, no difference in the time-series of superoxide is observed between the near cytosol and the MAM cytosol (Fig 4c).

### Simulation of Superoxide Effects on Calcium Dynamics

To simulate experimental observations indicating the influence of superoxide on calcium oscillations, the range of the superoxide production rate in the mitochondrial matrix (k_shunt_) was divided into two regions. The first region, analyzed initially, includes values of k_shunt_ for which, due to the higher degradation rate of superoxide in the cytosol compared to its leakage from a single mitochondrion, superoxide does not accumulate in the mitochondrion adjacent regions (i.e., MAM cytosol and near cytosol). As a result, superoxide oscillations driven by the ROS-induced ROS release (RIRR) mechanism (15) do not occur. Values in this region will hereafter be referred to as the non-oscillator value of k_shunt_, encompassing k_shunt_ values below 0.32 mM/s (Fig 4a).

#### Calcium Dynamics under Non-Oscillatory Superoxide

To investigate changes in calcium dynamics under the effect of redox-modulated IP3R channels, a bifurcation analysis of calcium was performed at a non-oscillator k_shunt_ (0.31 mM/s) under two scenarios: 1. With redox dynamics affecting calcium behavior through Eq 23 by modulating the value of K_P_ 2. With no influence of redox dynamics on IP3R function, i.e., with K_P_ held constant These two cases are presented in Fig 5.

**Fig 5:**
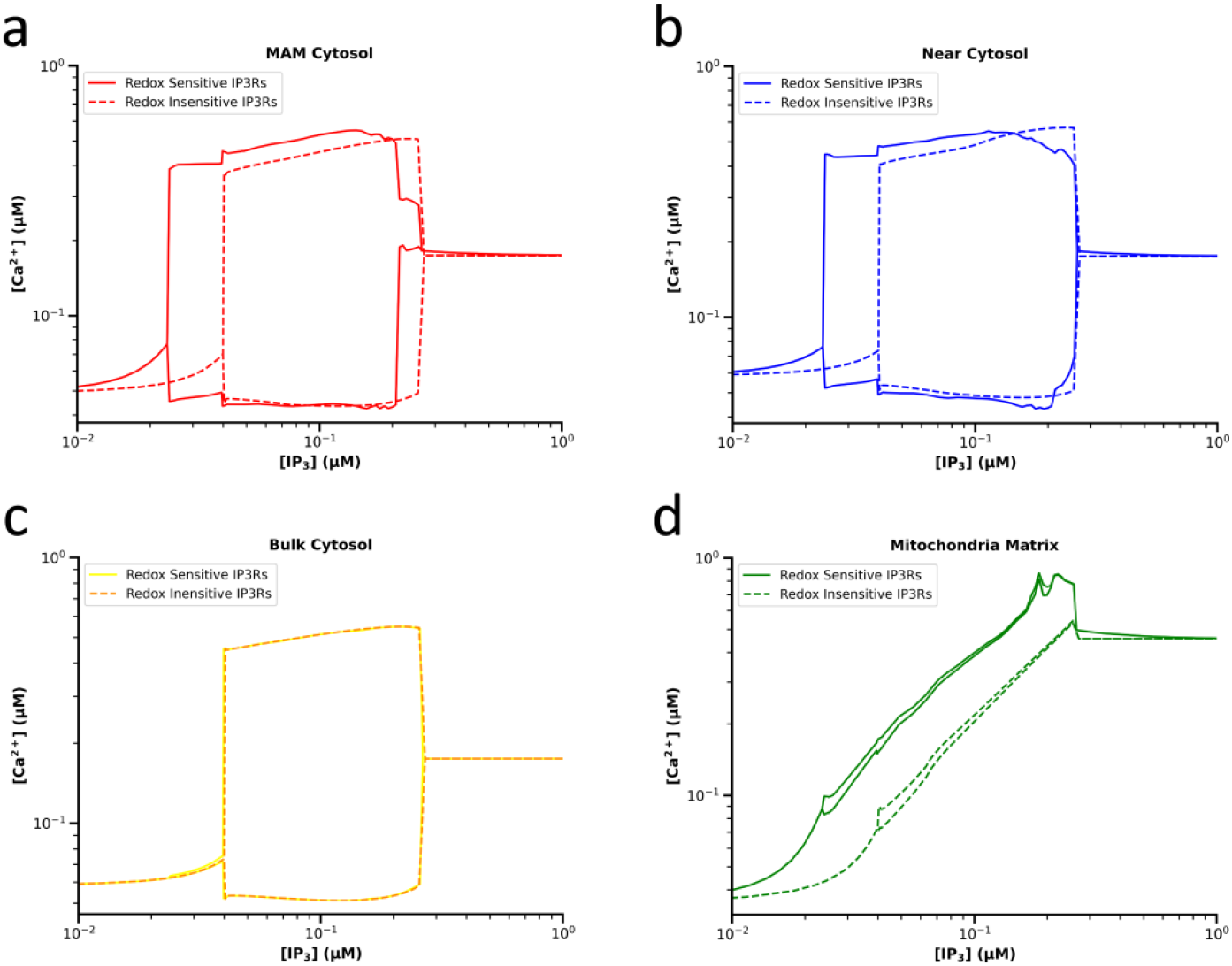
Bifurcation of calcium dynamics with and without the influence of redox state on IP3R function under non-oscillator k_shunt_. (a), (b), (c), And (d) show the calcium bifurcation based on IP3 concentration in the MAM cytosol, near cytosol, bulk cytosol, and mitochondrial matrix, respectively. Each panel compares redox-sensitive IP3Rs (solid curves) with redox-insensitive channels (dashed curves), where the parameter K_P_ is fixed at 0.2 µM.

Fig 5 shows that applying the oxidative effect of superoxide on calcium dynamics—via a reduction in K_P_ of IP3R channels, which represents the half-maximal concentration of IP3 for feedback on channel—leads to a leftward shift in the IP3 concentration required to initiate calcium oscillations in the MAM cytosol and near cytosol (Fig 5a and 5b). In contrast, there is no significant change in the bifurcation structure of calcium oscillations in the bulk cytosol (Fig 5c). Thus, under oxidative modulation of IP3R channels, basal concentrations of IP3 are sufficient to induce calcium oscillations in mitochondria-adjacent regions, whereas these concentrations are insufficient to trigger global calcium oscillations in the bulk cytosol. This behavior effectively simulates calcium flashes, sparks or puffs, which represent localized calcium oscillations within restricted cytosolic regions (7,10,19–21) (Fig 1a).

The absence of changes in calcium dynamics in the bulk cytosol following the application of redox effects to IP3R channels is due to the negligible oxidation of these channels compared to those faced to the MAM cytosol and near cytosol. This difference is itself due to the significantly lower superoxide concentration in the bulk cytosol relative to the mitochondria-adjacent regions (Fig 4a and 4d).

Fig 5d further demonstrates that the oxidative modulation of IP3R channels by superoxide leads to an increase in both the baseline and average calcium concentrations in the mitochondrial matrix. This outcome also reproduces experimental observations reporting elevated mitochondrial calcium levels under oxidative stress conditions (17,18).

Additionally, several studies have shown that redox condition of cell cytosol directly correlates with the number of calcium flashes over time (i.e., the frequency of calcium flashes). Specifically, increasing superoxide levels are associated with higher calcium flash frequencies (19,21), while scavenging superoxide using agents such as superoxide dismutase itself (19) or its mimetics (21) leads to a decrease in calcium flash frequency. To evaluate the model’s ability to replicate this behavior, we analyzed calcium dynamics under the vast range of non-oscillator k_shunt_ across a range of IP3 concentrations slightly broader than the bifurcation range shown in Fig 5a. We then computed the calcium oscillation frequency in various compartments of the model (Fig 6).

**Fig 6:**
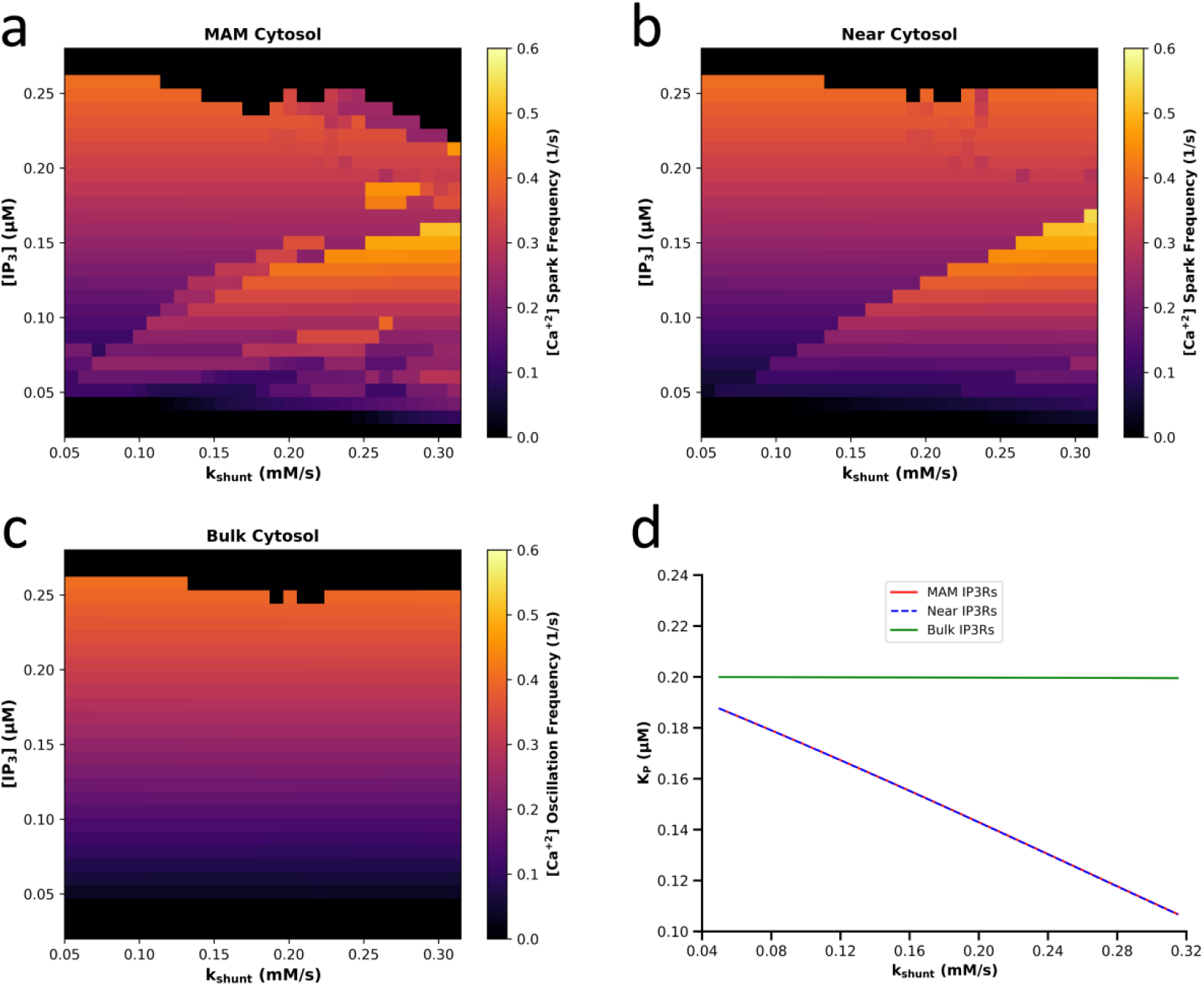
Calcium oscillation frequency at non-oscillator range of k_shunt_ and bifurcation range of IP3. (a), (b), And (c) respectively show the frequency of calcium oscillations in the MAM cytosol, the near cytosol, and the bulk cytosol. (d) Illustrates changes in the K_P_ parameter of IP3R channels as a function of superoxide concentration (driven by k_shunt_) in these three regions.

As expected, due to the minimal increase in superoxide levels in the bulk cytosol compared to the mitochondria-adjacent regions upon increasing the superoxide production rate (k_shunt_), the value of K_P_ in IP3R channels within the bulk cytoplasm remains nearly unchanged with increasing k_shunt_ (Fig 6d). In contrast, the K_P_ of IP3Rs in the MAM region and near cytoplasm decreases monotonically in response to increased channel oxidation, which is itself due to increased superoxide concentration in these regions. This decrease in K_P_ results in elevated sensitivity of IP3Rs to their stimulus, i.e., IP3 (see Eq S35, S37 and S40).

Consequently, this elevated sensitivity of IP3Rs to IP3 in channels adjacent to a mitochondrion, leads to a consistent increase in local calcium oscillation frequency in fixed IP3 levels (Fig 6a and 6b). Furthermore, the difference between the calcium frequency heatmaps for the mitochondria-adjacent regions and that of the bulk cytosol at basal IP3 concentrations (i.e., <0.04 μM) results from the distinct modulation of K_P_ in these regions. As a result, in low IP_3_ concentrations where the calcium in bulk cytosol does not oscillate, increased superoxide levels initially trigger localized calcium oscillations (or calcium flashes), which are then followed by a rise in their frequency. Thus, the model behavior shows good agreement with experimental observations regarding the effect of superoxide on local calcium dynamics at non-oscillator superoxide concentrations.

Irregularities in calcium dynamics within the bulk cytosol under very high IP3 concentrations (i.e., >0.25 μM, Fig 6a-6c) can be attributed to the inherent instability of calcium dynamics in the right side of the bifurcation region (based on IP3). In this condition, even slight, biologically insignificant perturbations profoundly impact oscillatory behavior of calcium. The same irregularities in calcium dynamics across the all three cytosolic compartments (Fig 6a-6c) further supports this explanation. Therefore, such high IP3 concentrations do not appear to have physiological significance.

It is also important to note that in IP3 concentrations where calcium oscillations occur not only in the MAM cytosol and near cytosol but also in the bulk cytosol, if the calcium peaks in the bulk cytosol closely align in time with those in the other two regions (as shown Fig 3d), then distinguishing localized calcium flashes from global calcium oscillations in fluorescence microscopy of cells becomes practically infeasible.

In general, modeling the modulation of IP3R gating dynamics under the influence of superoxide—rather than other oxidants—is supported by two key rationales: First, among the thiol-oxidizing agents whose effects on calcium dynamics via IP3Rs have been studied—such as thimerosal (30,51), diamide (27), superoxide (26), peroxide (19), vanadate (29), chloramine T (29), etc.—only superoxide and peroxide are naturally occurring compounds endogenously produced within cells, particularly by mitochondria. Second, the diffusion coefficient of peroxide in aqueous solutions has been reported to be 10 to 100 times greater than that of superoxide (54). Therefore, given the highly localized nature of calcium flashes (typically within a radius of 1–2 micrometers), modeling a superoxide-mediated effect is more plausible and physiologically consistent than one mediated by peroxide.

#### Calcium Dynamics Under Oscillatory Superoxide

Another category of calcium oscillation observations, especially calcium flashes, has been conducted under conditions in which not only calcium but also reactive oxygen species (ROS), including superoxide, exhibit oscillatory behavior. For instance, Zhou et al. (20) carefully investigated the frequency changes of calcium flashes synchronized with high-amplitude oscillations of the mitochondrial membrane potential, attributed to the opening of IMAC channels. They quantified that the frequency of calcium flashes during the opening of IMAC channels—that are the primary pathway for superoxide flux from the mitochondrial matrix to the intermembrane space and cytosol—is on average 2.5 times greater than that observed when the IMAC channels are closed.

Accordingly, to simulate this phenomenon, we examined the higher range of k_shunt_, which corresponds to the state wherein, due to elevated mitochondrial superoxide production rate, superoxide accumulates outside the mitochondria and, through positive feedback on IMAC channels, induces superoxide oscillations (i.e., ROS-induced ROS Release). In consequence, based on our model for the redox dynamics of IP3R channels influenced by superoxide (Eq 24-28), it was expected that superoxide oscillations would also modulate the frequency of calcium oscillations.

As a first step of this investigation, we studied the model behavior for a combination of the oscillator values of k_shunt_ and IP3. Fig 7 illustrates the oscillations of calcium, superoxide, and the parameter K_P_ of IP3R channels (affected by redox oscillations of the channels) for a specific oscillator value of k_shunt_ and a basal concentration of IP3 (a resting cellular state).

**Fig 7:**
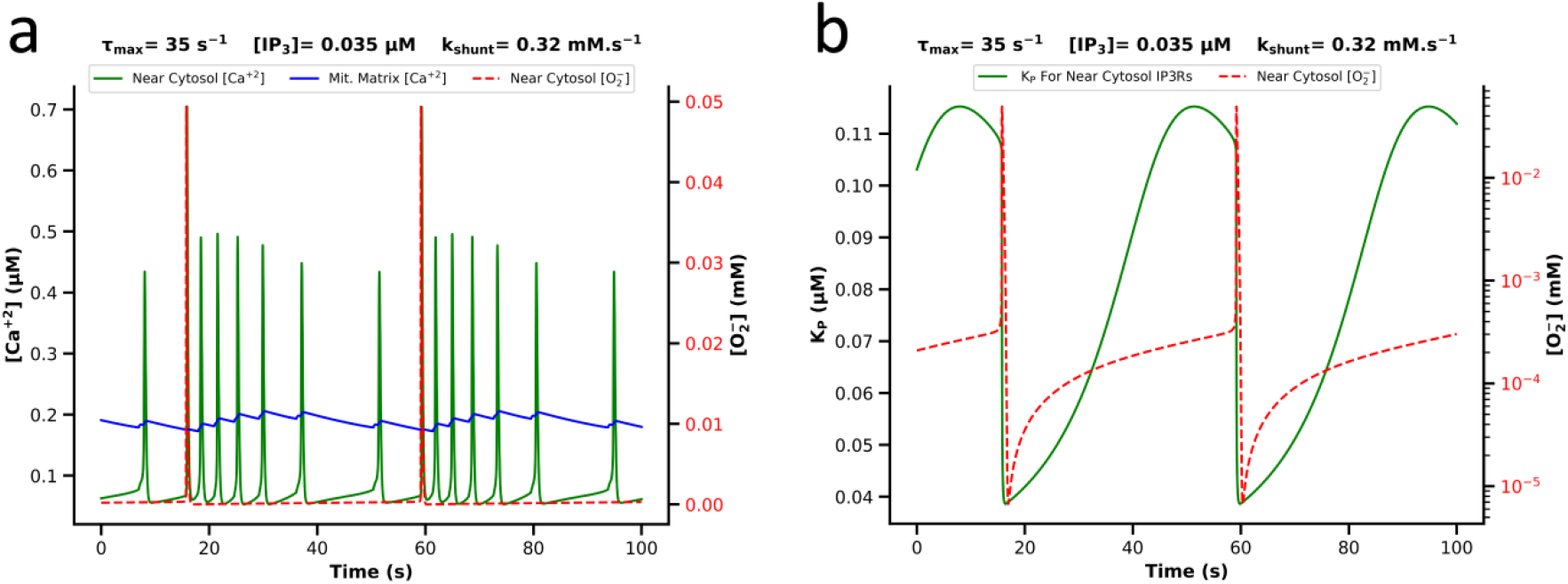
Oscillations of Superoxide, Calcium and K_P_ of IP3Rs. (a) The time-series of superoxide (red dashed curve) and calcium (green curve) in the near cytosol, and calcium in the mitochondrial matrix (blue curve). (b) The oscillation of superoxide in the near cytosol (red dashed curve) alongside the oscillation of K_P_ parameter of IP3R channels, mediated by redox state oscillation. Both graphs are plotted for τ_max_= 35 s^−1^ and a specific k_shunt_ and IP3, indicated above the graphs.

Fig 7b clearly illustrates how, during each superoxide pulse from a mitochondrion, the adjacent IP3R channels undergo oxidation, resulting in a decrease in their K_P_ according to Eq 23. Once the IMAC channels close and the oxidation of IP3R channels subsides, K_P_ increases again. Fig 7a demonstrates the consequences of this oscillation of K_P_ in near cytosol calcium dynamics, which is through altering the open probability dynamics of the IP3Rs (see Eq S53, S55 and S58), thereby modulating the frequency of calcium flashes. To gain a more intuitive understanding of how superoxide pulses can influence calcium flashes— as observed under confocal microscopy (20)—see Supplementary Video.

As the second step, to investigate the ratio of calcium oscillation frequency during IMACs opening to that during IMACs closure, we re-solved the model for each combination of k_shunt_ and IP3. This time, however, the K_P_ value of the IP3Rs was held constant at its maximum value observed during superoxide oscillations in the first step, in order to simulate calcium dynamics under the condition where IMAC channels remain closed and superoxide does not oscillate. It is important to note that the maximum K_P_ value varies under different mitochondrial superoxide production rates (i.e., for each k_shunt_). Fig 8 represents the calcium dynamics under both oscillatory K_P_ (first step) and fixed K_P_ (second step) conditions for a specific k_shunt_ and IP3.

**Fig 8:**
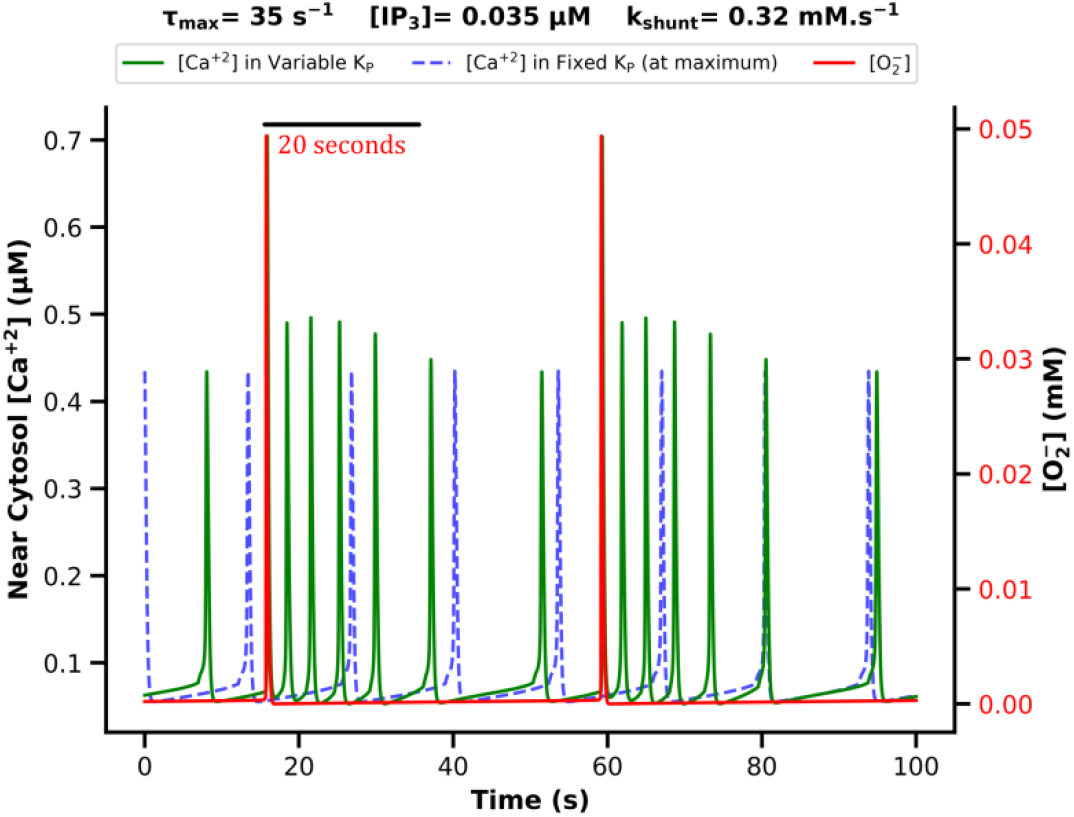
Oscillations of superoxide (red curve) and calcium under variable K_P_ (green curve), and fixed K_P_ (blue dashed curve). This figure presents the simulation results for a specific k_shunt_ and IP3. The highlighted 20-second time interval marks the time window during which the number of calcium flashes under variable K_P_ was counted following each superoxide pulse (for a description of its use, see the following text).

Based on the findings of Zhou et al. (20), which indicate that the increased frequency of calcium oscillations during each IMACs opening and associated superoxide pulse lasts approximately 20 seconds, we calculated the frequency of calcium oscillations only within the first 20 seconds following each superoxide pulse under the variable K_P_ condition. This was then compared to the frequency observed under fixed K_P_ condition (i.e. fixed at its maximum for each k_shunt_) resulting in a similar ratio to that calculated by Zhou et al. (20). This ratio, hereafter referred to as the calcium frequency modulation ratio, quantifies the change of calcium oscillation frequency under the oscillatory dynamics of superoxide. The convergence between the calcium time-series under variable K_P_ (green curve in Fig 8) and fixed K_P_ (blue dashed curve in Fig 8) increases as this ratio approaches 1. Fig 9 shows this ratio in simulations performed over the range of oscillator k_shunt_ and bifurcation range of IP3.

**Fig 9:**
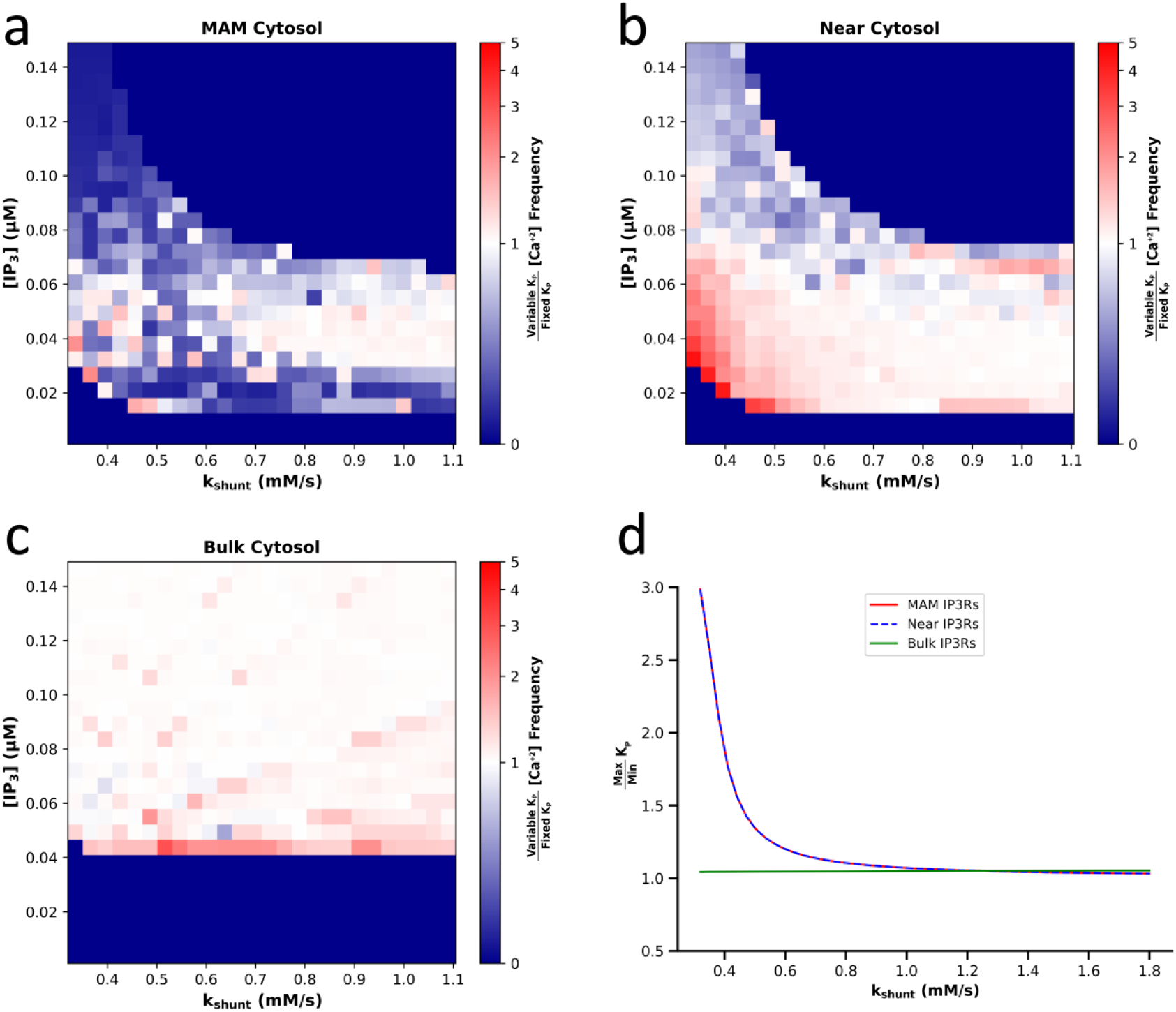
The calcium frequency modulation ratio. (a), (b) And (c) respectively shows the calcium frequency modulation ratio—defined as the ratio of calcium oscillation frequency under variable K_P_ to fixed K_P_—in the MAM cytosol, near cytosol, and bulk cytosol, across different values of k_shunt_ and IP3. (d) Shows the ratio of the maximum to minimum K_P_ for each specific k_shunt_ value.

Comparison of Fig 9a-9c indicates that the most significant and regular shift in the calcium frequency modulation ratio by altering IP3 and k_shunt_ occurs in the near cytosol. Indeed, as expected, the calcium dynamics in the large volume of the bulk cytosol—similarly to the non-oscillatory superoxide condition where changes in superoxide production rate of a single mitochondrion did not affect bulk calcium dynamics—is also insensitive to variations in the superoxide oscillation frequency (Fig 9c). On the other hand, the irregular shifts in the calcium frequency modulation ratio in the MAM cytosol under oscillatory superoxide conditions is mainly due to the high instability of calcium oscillations and fundamental alterations in the shape of calcium pulses induced by K_P_ oscillation in each superoxide pulse in this area (see Fig S8). This high instability itself, arises from the extremely limited volume of the MAM cytosol compared to the bulk and near cytosol regions, which leads to extended peaks in calcium concentration and IP3R open probability, thereby significantly disrupting regular trends in this compartment.

The most critical result of this section is Fig 9b, which shows that in the lower-left region of the heatmap— i.e., at low IP3 concentrations (0.01 to 0.04 µM) and in the initial range of oscillator k_shunt_ values (less than 0.06 mM.s^-1^)— the calcium frequency modulation ratio is greater than 1 and reaches values as high as 5. Fig 9b also shows that at these low IP3 concentrations, increasing k_shunt_ (i.e., mitochondrial superoxide production rate) leads to a decrease in this modulation ratio. This is because as k_shunt_ increases—leading to a higher frequency of superoxide oscillations (Fig 4b)— the recovery time for the redox state of IP3Rs before the next superoxide pulse decreases. Consequently, the ratio of the maximum to minimum IP3R_RED_ (i.e., IP3Rs reduction fraction in Eq 22)—and thus the ratio of maximum to minimum K_P_ of IP3Rs— decreases and approaches 1 (Fig 9d). Accordingly, when the cellular reductive capacity remains constant, increasing superoxide oscillation frequency diminishes its modulatory effect on calcium flash frequency, as shown in Fig 9b. Insufficient time for K_P_ to increase again at high k_shunt_ also results in IP3R channels remaining fully open at high IP3 levels, explaining the absence of calcium oscillations in the upper-right region of the heatmaps for both MAM cytosol (Fig 9a) and near cytosol (Fig 9b).

It is pertinent to note that the current model for the redox regulation of IP3R channels (i.e., Eq. 23) is predicated on the quasi-linear assumption that alterations in the K_p_ variable—a key gating controller of IP3R channels—occur non-cooperatively. Consequently, a hyperbolic function, rather than a sigmoidal one, was employed for this formulation (i.e., Eq 23). In the absence of precise biophysical data elucidating the underlying biophysical nature of the K_p_ parameter and its exact dynamic response to redox fluctuations, this assumption strictly adheres to the principle of parsimony in scientific modeling, provided it successfully reproduces the desired macroscopic behavior. Nevertheless, a hallmark of (positively) cooperative models utilizing a sigmoidal (Hill) function is their capacity to confer a switch-like dynamic upon the K_p_ parameter—a characteristic underlying a significant portion of critical cellular behaviors (55,56). This fundamental difference in K_p_ modulation is illustrated in Fig S7.

Next, to investigate the physiological significance of these changes in calcium oscillations, we examined the effect of varying oscillator k_shunt_ and IP3 levels on the average calcium concentration within the mitochondrial matrix (Fig 10). First, we evaluated the average calcium concentration in the mitochondrial matrix with normal IP3Rs that K_P_ is variable and oscillates due to the redox oscillation of channels (Fig 10a). Then, to assess strictly the effect of oscillatory superoxide on mitochondrial calcium, we calculated the ratio of average mitochondrial calcium concentration under variable (oscillatory) K_P_ to that under fixed K_P_ (at its maximum in each k_shunt_) conditions (Fig 10b).

**Fig 10:**
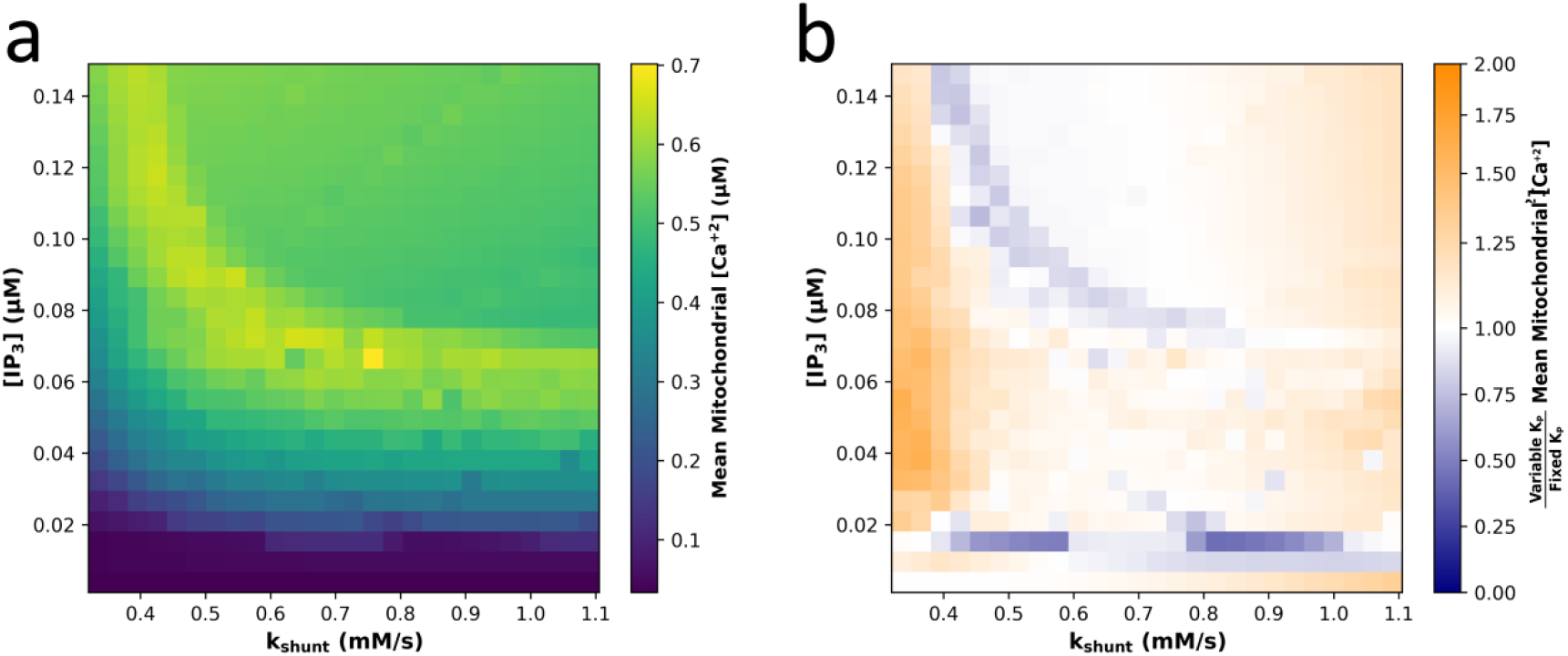
The average mitochondrial calcium concentration under varying superoxide production rate and IP3 levels. (a) Illustrates the mean calcium concentration within the mitochondrial matrix at different values of k_shunt_ and IP3. (b) Shows the ratio of average mitochondrial calcium concentration under oscillatory K_p_ (variable) versus non-oscillatory K_p_ (fixed at its maximum) conditions.

Fig 10a demonstrates that, at each k_shunt_, increasing IP3 raises the average mitochondrial calcium concentration as long as calcium continues to oscillate within the MAM cytosol. However, when IP3 level becomes excessively high—leading to sustained opening of IP3R channels and termination of calcium oscillations in both the near cytosol and MAM cytosol—the average mitochondrial calcium concentration drops. This result is fully consistent with the known properties of mitochondrial calcium uniporters (MCU), which are low-affinity, high-capacity calcium transporters (57,58) and thus, function most effectively at peaks of calcium oscillations in the MAM cytosol and near cytosol.

Additionally, Fig 10a indicates that at low IP3 concentrations—where cytosolic calcium oscillations appear only as localized flashes in the mitochondria-adjacent cytosol—increasing the mitochondrial superoxide production rate initially increases and then stabilizes the average mitochondrial calcium concentration. This observation aligns with the reduced calcium frequency modulation ratio at high superoxide oscillation frequencies.

As anticipated, Fig 10b reveals that mitochondrial calcium dynamics, like those in the cytosol (i.e., calcium flashes), are influenced by oscillatory dynamics of superoxide only at low frequencies of superoxide oscillation. As the superoxide oscillation frequency increases, the impact of its oscillatory dynamics converges its steady state effect on the average mitochondrial calcium concentration.

In the final part of the simulation of experimental studies, we examined the model’s behavior across different values of the parameter τ_max_, which, according to the model by Sneyd et al. (44), determines the intrinsic calcium oscillation frequency in various cell types. We explored different values of τ_max_ within the oscillatory range of k_shunt_ and low IP3 concentration (0.03 μM) to focus on the behavior of localized calcium flashes. Fig 11 displays the ratio of calcium oscillation frequency under variable K_P_ to that under fixed K_P_ conditions.

**Fig 11:**
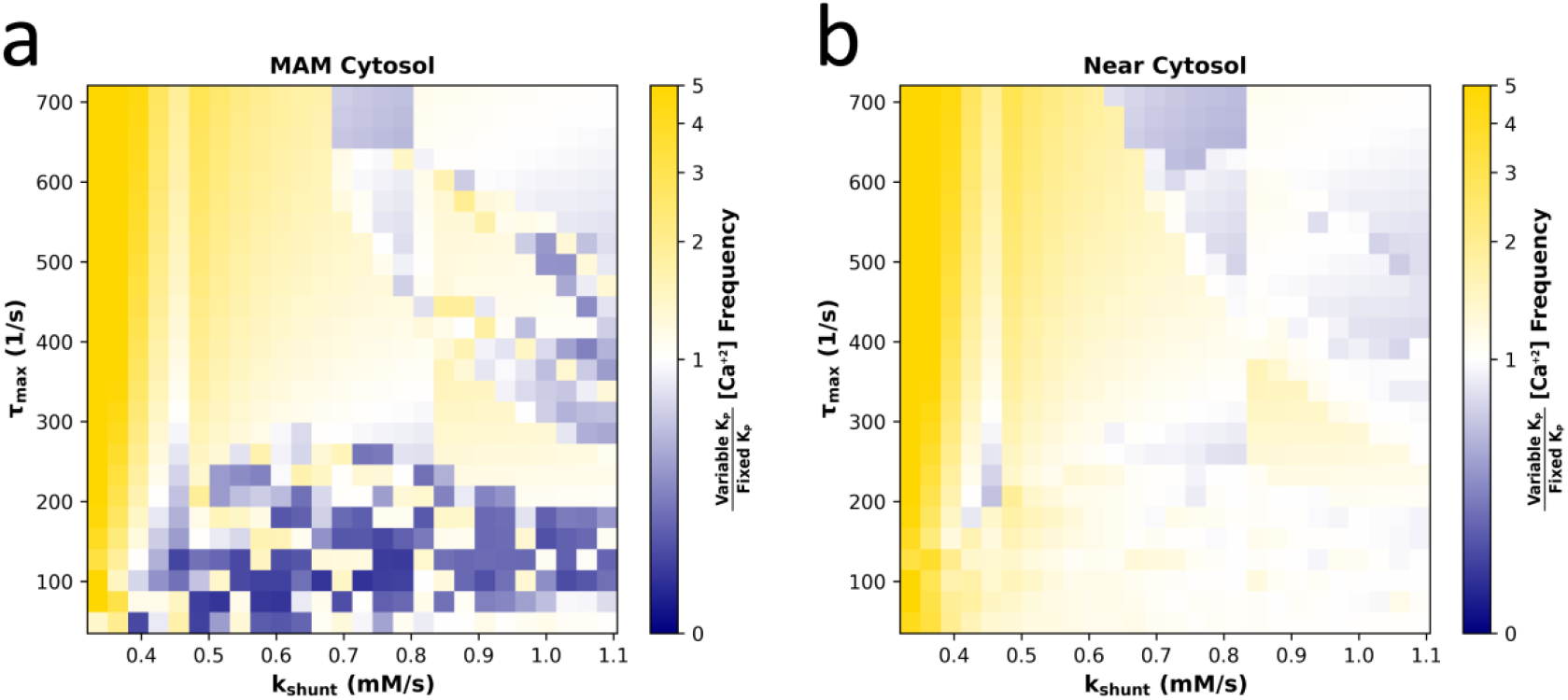
The calcium frequency modulation ratio across different values of τ_max_ and k_shunt_. (a) Shows this ratio in the MAM cytosol and (b) presents it in the near cytosol.

Both Fig 11a and 11b again demonstrate that oscillatory dynamics of superoxide is physiologically significant only at low rates of superoxide production that result in low frequencies of superoxide oscillations. As this rate (k_shunt_) increases, the effect of superoxide oscillations on calcium dynamics becomes less distinct. Ultimately, Fig 11 indicates that the current model is capable of simulating the impact of oscillatory dynamics of superoxide on calcium flashes—as reported in experimental observations— across a wide range of non-excitable cell types with varying intrinsic calcium oscillation frequencies.

### Model Predictions

Numerous studies have reported that the calcium dynamics in cytosol and within the mitochondrial matrix are altered by changes in the length of the MAM space (59). Specifically, with an increase in the MAM length and enhanced connectivity between the mitochondria and the endoplasmic reticulum, the average mitochondrial calcium level rises (60). Conversely, a reduction in MAM length leads to insufficient calcium uptake by the mitochondria, impairing ATP production (48).

Therefore, to examine the geometric significance of the MAM cytosol in the current model and to predict the model’s behavior under varying MAM lengths, we solved the model for different MAM lengths at a resting IP3 concentration (in order to focus on calcium flashes). This analysis involved evaluating the frequency of calcium flashes and the average mitochondrial calcium concentration across a range of MAM lengths. The results are presented in Fig 12.

**Fig 12:**
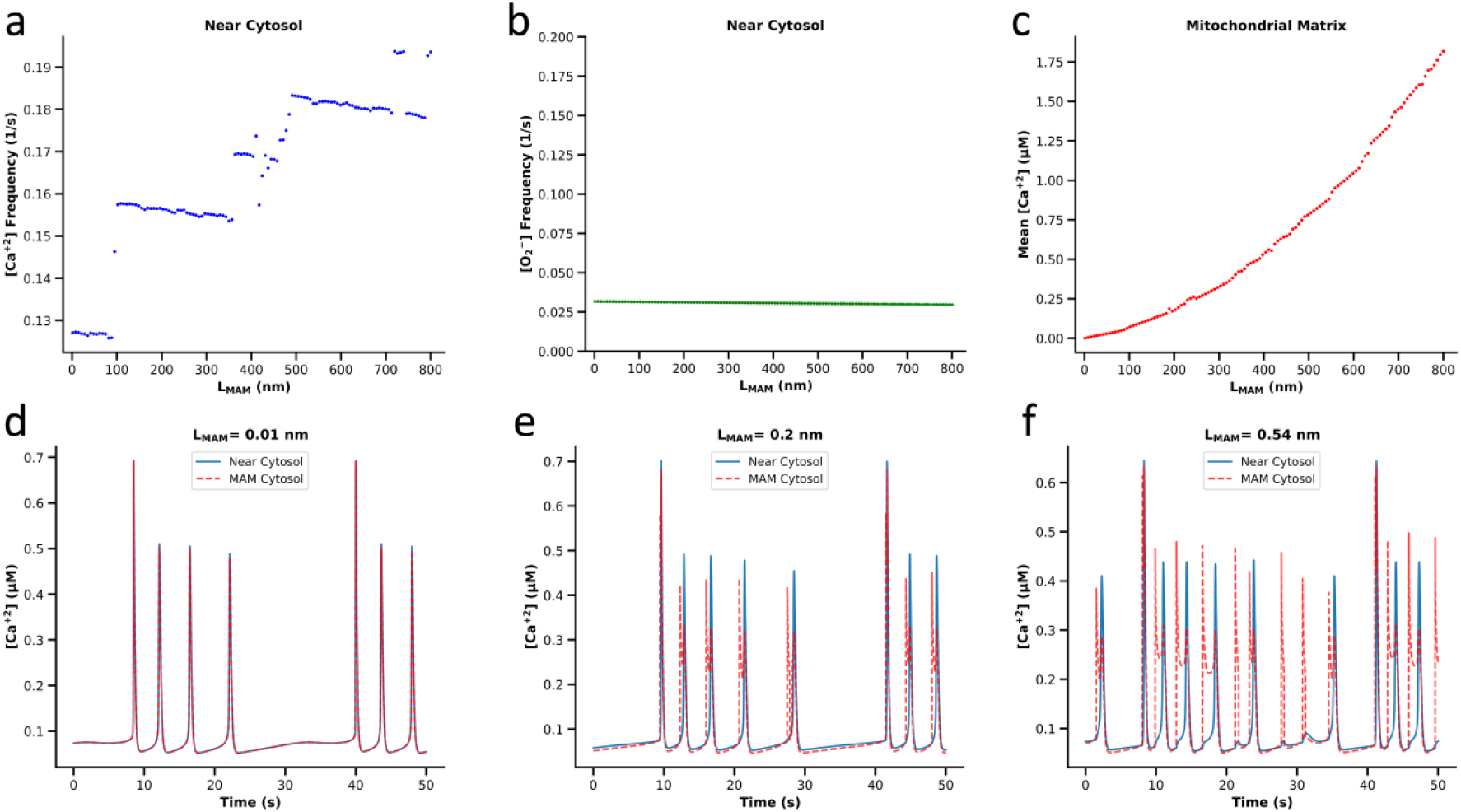
Effect of MAM Length on Superoxide and Calcium Dynamics. (a) And (b) respectively illustrate the frequency of calcium flashes and superoxide flashes in the near cytosol as a function of MAM length (each point represents the simulation result at a specific MAM length). (c) Shows the average mitochondrial calcium concentration with changing MAM length. (d), (e) And (f) depict calcium oscillations in the MAM cytosol (red dashed curve) and near cytosol (blue curve) for three different MAM lengths.

Interestingly, Fig 12a indicates that the frequency of calcium oscillations in the near cytosol (calcium flashes) follows a rather direct, albeit stepwise, proportional relationship with MAM length. This stepwise behavior arises from the fact that the primary regulator of calcium dynamics is superoxide oscillations, which manifests as very high-amplitude peaks, clearly visible in Fig 12d-12f, and thus the discrete increase in the number of calcium flashes per unit time occurring between two superoxide pulses leads to a sudden steplike rise in the mean frequency of calcium flashes. This can be observed in the calcium dynamics at the three different MAM lengths shown in Fig 12d-12f.

To understand the underlying reason for the increase in calcium flash frequency with MAM length, we also plotted the frequency of superoxide flashes for each MAM length (Fig 12b). This analysis clearly demonstrates that, unlike calcium, superoxide oscillation frequency remains unaffected by changes in MAM length. Therefore, the observed variation in calcium flash frequency with MAM length is not due to changes in superoxide dynamics.

Indeed, one effect of increasing the volume of the MAM cytosol is that, as this volume expands, the amount of calcium diffusing from the MAM cytosol to the near cytosol during each calcium pulse increases. This, in turn, causes the IP3Rs in the near cytosol to reach their calcium-based activation threshold more quickly. This mechanism explains why an increase in MAM space volume leads to a higher frequency of calcium flashes in the near cytosol. At last, consistent with experimental observations (61), Fig 12c shows a direct relationship between MAM length and the average mitochondrial calcium concentration.

### Sensitivity Analysis

Given that the majority of parameter values are derived from four distinct, previously developed models, concerns may arise regarding the biological compatibility and physiological plausibility of this complex model and parameter integration. However, two of the reference models employed in the present study— namely, the calcium dynamics model by Sneyd et al. (44) and the model by Moshkforoush et al. (38)— serve as generic frameworks applicable to a broad spectrum of non-excitable cells. Furthermore, the model proposed by Wacquier et al. (36) was specifically formulated for HeLa cells, which are widely regarded as typical non-excitable cells in the context of calcium dynamics. Moreover, although the model by Yang et al. (13) was initially developed to describe mitochondrial reactive oxygen species (ROS) dynamics in excitable cardiac cells, it has previously been successfully applied to non-excitable cancer cells conserving the most of original parameter set (62), demonstrating a high degree of consistency in simulating ROS dynamics within these cells.

Nonetheless, to rigorously demonstrate the high biological robustness of the proposed model and its applicability across diverse cell types and physiological contexts, we conducted a two-stage hierarchical Global Sensitivity Analysis (GSA) based on the Regional Sensitivity Analysis (RSA) framework, and Monte Carlo Filtering (MCF). The first-stage aimed to establish that local calcium flashes—which are critical for maintaining essential cellular energetics— and mitochondrial ROS oscillations can be robustly generated across a broad range of the parameter space in the present model. To this end, a ±40% variation range was assigned to each parameter, and 100,000 distinct parameter sets were sampled from the multidimensional space using Latin Hypercube Sampling (LHS). Following the simulation of all generated samples, those exhibiting physiological local calcium oscillations in the near cytosol (defined by a basal calcium level of < 0.2 μM and a peak-to-trough ratio of > 1.8), alongside ROS oscillations in the near cytosol (defined by a peak-to-trough ratio of > 10), were classified as favorable (behavioral) states; whereas the remainder were categorized as unfavorable (non-behavioral) states. Fig 14 presents the Kolmogorov-Smirnov (KS) statistics for the relatively important parameters, derived from their cumulative distribution functions (CDFs) under defined favorable and unfavorable conditions (Note: Only parameters yielding a KS value greater than 0.1 are considered relatively important and depicted here; the CDF profiles and corresponding KS values for the entire parameter set are provided in Fig S10,S11).

As evident from Fig 13 (and Fig S10,S11), the low KS values (<0.3) indicate that none of the parameters in the developed model are restricted to a narrow functional range for generating local ROS and calcium oscillations. It is also noteworthy that during this stage of the GSA, approximately 39% of the 100,000 generated parameter sets successfully yielded the defined behavioral state. Consequently, this clearly demonstrates that despite being synthesized from a complex integration of multiple pre-existing models, the present framework robustly preserves biological consistency. Furthermore, Fig 13 demonstrates that the most important parameters determining the activation/deactivation of local calcium and ROS oscillations in our model fall into four categories: a geometrical parameter (i.e., R_mit_), parameters linked to the intrinsic dynamics of IP3R channels (i.e., K_h_ and K_C_), those related to the ROS dynamics (i.e., h_IMAC_ and D_S_), and parameters controlling the redox dynamics of IP3R channels (i.e., g_max_ and k_RED_).

**Fig 13:**
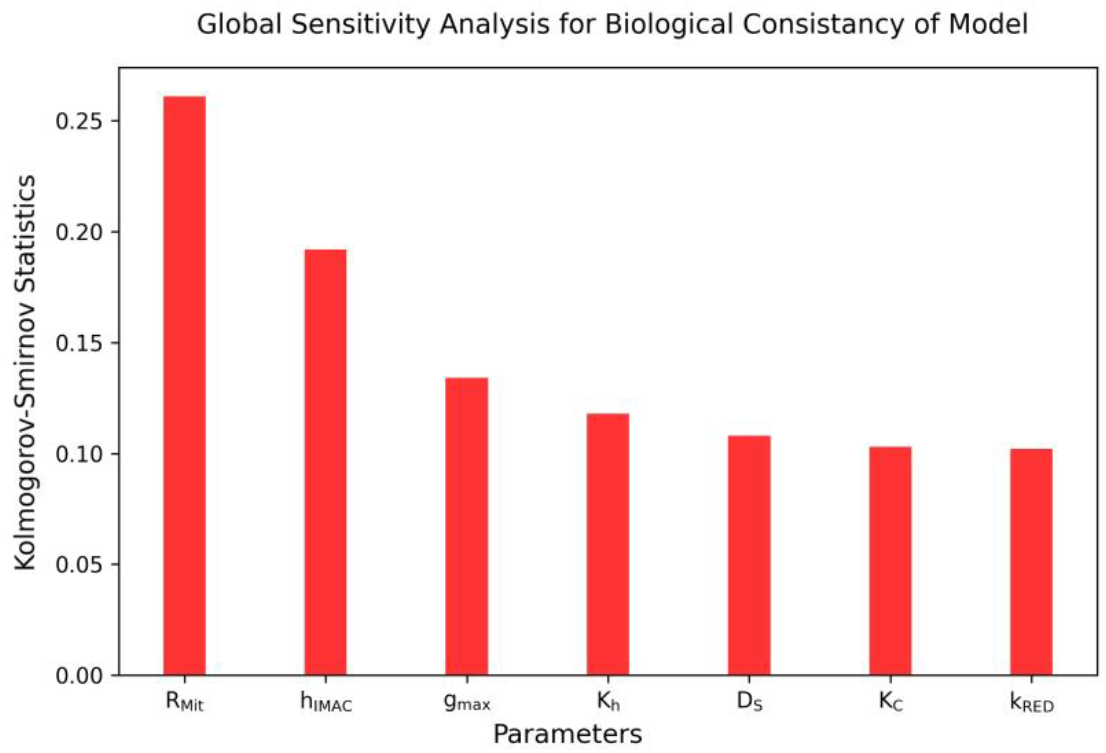
First-stage global sensitivity analysis of model parameters. The figure illustrates the Kolmogorov-Smirnov (KS) statistics derived from the comparison of cumulative distribution functions (CDFs) between favorable (oscillatory calcium in the Near cytosol) and unfavorable (non-oscillatory calcium) states. Only parameters exhibiting a KS statistic greater than 0.1 are displayed.

In the second-stage of the analysis, to verify that the proposed model avoids overtuning and can robustly reconstruct ROS-mediated calcium frequency modulation across a broad range for each of its 52 parameters—rather than relying on specific, fine-tuned values—100,000 parameter combinations (sampling by LHS with a ±40% variation range) were simulated under two distinct conditions: ROS-calcium coupled dynamics and uncoupled state. In this stage, favorable (behavioral) outcomes were explicitly defined as instances where, alongside the maintenance of both ROS and calcium oscillations, the calcium oscillation frequency exhibited a significant absolute shift of greater than 20% in the coupled state compared to the uncoupled one.

Although a mere 0.034% (n=34) of the sampled combinations were classified as favorable, the CDF plots in Fig S12,S13 illustrate that each parameter individually can, in principle, retain a wide permissible range to satisfy these criteria. Interestingly, this exceedingly low percentage of favorable states highlights a crucial biological safeguard: independent variations in individual parameters are generally insufficient to alter the calcium oscillation frequency, which serves as a vital cellular signaling tool. This inherent robustness protects frequency-sensitive signaling pathways (11) from random, disruptive fluctuations. Ultimately, this statistical phenomenon reflects the Curse of Dimensionality (63), demonstrating a profound interdependence among the model parameters to execute such a meaningful shift in calcium frequency. To illustrate this mathematically, assuming a hypothetical scenario of independent parameters, if each of the 52 parameters could yield the favorable behavior of the global model across 85% (0.85) of its respective range, the probability of a randomly generated sample having all 52 parameters simultaneously fall within their optimal bounds would be 0.85 to the power of 52, which equates to approximately 0.021%.

Finally, to explicitly quantify the direction and magnitude of the calcium dynamics response to minor perturbations (5%) specifically within the newly proposed IP3R redox model, a linear sensitivity analysis was performed for each of 5 newly proposed parameters. The results of this analysis are summarized in Table 1.

**Table 1:** Linear sensitivity analysis of parameters the proposed IP3R redox regulation model. C is the amplitude of calcium oscillation in different compartments.

| | $\frac{\% \Delta C_{MAM}}{\% \Delta Parameter}$ | $\frac{\% \Delta C_{Near}}{\% \Delta Parameter}$ | $\frac{\% \Delta C_{Mit}}{\% \Delta Parameter}$ |
| --- | --- | --- | --- |
| **k<sub>OX</sub>** | 1.767 | -0.115 | 2.223 |
| **k<sub>RED</sub>** | -1.071 | 1.711 | -2.146 |
| **K<sub>IP3R, OX</sub>** | -0.32 | -0.395 | 0.775 |
| **g<sub>max</sub>** | 0.266 | 0.302 | 4.272 |
| **G<sub>OX</sub>** | -0.062 | -0.075 | -0.965 |

## Conclusion

The mathematical model developed in this study successfully captured and simulated the influence of superoxide dynamics on local calcium oscillations as observed in experimental studies. Compared to previous models addressing the impact of reactive oxygen species (ROS) on calcium dynamics, our approach adopts a more rational framework by incorporating the influence of ROS through modulation of the redox state governing IP3 receptors (IP3R) gating dynamics, rather than assuming a direct and phenomenological effect of ROS on these calcium channels. This highlights how existing models of IP3R dynamics can be extended to account for redox-dependent modifications under varying cellular conditions.

Nonetheless, the current model has two limitations. First, it considers only the unidirectional interaction between ROS and calcium—specifically from mitochondrial superoxide to IP3R-mediated calcium dynamics—whereas this relationship is inherently bidirectional. The second is that the model contains only the interactions between calcium and superoxide in the vicinity of a single mitochondrion, without accounting for the role of other mitochondria as part of the broader mitochondrial network.

Therefore, integrating the bidirectional relationship of ROS and calcium and the structural and functional dynamics of the mitochondrial network into the proposed model in future studies, can provide a more comprehensive and physiologically realistic representation of calcium and ROS signaling under both normal and pathological cellular conditions.

Ultimately, beyond elucidating fundamental biophysical interactions, the proposed framework holds significant practical applications for advancing both basic and translational research. First, by establishing a mechanistic link between localized oxidative stress and calcium frequency decoding, the model provides a crucial computational tool for investigating pathological conditions characterized by altered redox homeostasis, such as neurodegenerative diseases and aging. Such predictive capabilities can significantly accelerate hypothesis generation, minimize trial-and-error, and guide future experimental and therapeutic designs.

## Acknowledgments

We are grateful to Professor James Sneyd for his valuable comments on an earlier draft of the manuscript. His suggestions, particularly regarding the justification of the model assumptions, helped improve the clarity and rigor of the presentation.

## References

1. Berridge MJ, Lipp P, Bootman MD. The versatility and universality of calcium signalling. Nat Rev Mol Cell Biol. 2000 Oct;1(1):11–21. doi:10.1038/35036035

2. Berridge MJ. The Inositol Trisphosphate/Calcium Signaling Pathway in Health and Disease. Physiological Reviews. 2016 Oct;96(4):1261–96. doi:10.1152/physrev.00006.2016

3. Dupont G, Combettes L, Bird GS, Putney JW. Calcium Oscillations. Cold Spring Harbor Perspectives in Biology. 2011 Mar 1;3(3):a004226–a004226. doi:10.1101/cshperspect.a004226

4. Jouaville LS, Ichas F, Holmuhamedov EL, Camacho P, Lechleiter JD. Synchronization of calcium waves by mitochondrial substrates in Xenopus laevis oocytes. Nature. 1995 Oct;377(6548):6548. doi:10.1038/377438a0

5. Tengholm A, Gylfe E. Oscillatory control of insulin secretion. Molecular and Cellular Endocrinology. 2009 Jan 15; Special Issue: Endocrine Aspects of Type II Diabetes 297(1):58–72. doi:10.1016/j.mce.2008.07.009

6. Thorn P, Lawrie AM, Smith PM, Gallacher DV, Petersen OH. Ca2+ oscillations in pancreatic acinar cells: spatiotemporal relationships and functional implications. Cell Calcium. 1993 Nov;14(10):746–57. doi:10.1016/0143-4160(93)90100-k PubMed PMID: 8131191.

7. Li Y, Zhang Z, Hao X, Yu T, Li S. Qiliqiangxin Capsule Modulates Calcium Transients and Calcium Sparks in Human Induced Pluripotent Stem Cell-Derived Cardiomyocytes. Evidence-Based Complementary and Alternative Medicine. 2022 Aug 30;2022:e9361077. doi:10.1155/2022/9361077

8. Lohmann C, Myhr KL, Wong ROL. Transmitter-evoked local calcium release stabilizes developing dendrites. Nature. 2002 Jul;418(6894):177–81. doi:10.1038/nature00850

9. Lohmann C, Finski A, Bonhoeffer T. Local calcium transients regulate the spontaneous motility of dendritic filopodia. Nat Neurosci. 2005 Mar;8(3):305–12. doi:10.1038/nn1406

10. Cárdenas C, Miller RA, Smith I, Bui T, Molgó J, Müller M, et al. Essential Regulation of Cell Bioenergetics by Constitutive InsP3 Receptor Ca2+ Transfer to Mitochondria. Cell. 2010 Jul;142(2):270–83. doi:10.1016/j.cell.2010.06.007

11. Smedler E, Uhlén P. Frequency decoding of calcium oscillations. Biochimica et Biophysica Acta (BBA) - General Subjects. 2014 Mar;1840(3):964–9. doi:10.1016/j.bbagen.2013.11.015

12. Ohadi D, Schmitt DL, Calabrese B, Halpain S, Zhang J, Rangamani P. Computational Modeling Reveals Frequency Modulation of Calcium-cAMP/PKA Pathway in Dendritic Spines. Biophysical Journal. 2019 Nov;117(10):1963–80. doi:10.1016/j.bpj.2019.10.003

13. Yang L, Korge P, Weiss JN, Qu Z. Mitochondrial Oscillations and Waves in Cardiac Myocytes: Insights from Computational Models. Biophysical Journal. 2010 Apr 21;98(8):1428–38. doi:10.1016/j.bpj.2009.12.4300

14. Park J, Lee J, Choi C. Mitochondrial Network Determines Intracellular ROS Dynamics and Sensitivity to Oxidative Stress through Switching Inter-Mitochondrial Messengers. PLOS ONE. 2011 Aug 4;6(8):e23211. doi:10.1371/journal.pone.0023211

15. Zorov DB, Juhaszova M, Sollott SJ. Mitochondrial Reactive Oxygen Species (ROS) and ROS-Induced ROS Release. Physiological Reviews. 2014 Jul;94(3):909–50. doi:10.1152/physrev.00026.2013

16. Wang W, Fang H, Groom L, Cheng A, Zhang W, Liu J, et al. Superoxide Flashes in Single Mitochondria. Cell. 2008 Jul;134(2):279–90. doi:10.1016/j.cell.2008.06.017

17. Görlach A, Bertram K, Hudecova S, Krizanova O. Calcium and ROS: A mutual interplay. Redox Biology. 2015 Dec;6:260–71. doi:10.1016/j.redox.2015.08.010

18. Brookes PS, Yoon Y, Robotham JL, Anders MW, Sheu SS. Calcium, ATP, and ROS: a mitochondrial love-hate triangle. American Journal of Physiology-Cell Physiology. 2004 Oct;287(4):C817–33. doi:10.1152/ajpcell.00139.2004

19. Isaeva EV, Shkryl VM, Shirokova N. Mitochondrial redox state and Ca ^2+^ sparks in permeabilized mammalian skeletal muscle: Mitochondrial redox potential and Ca ^2+^ sparks in muscle. The Journal of Physiology. 2005 Jun 15;565(3):855–72. doi:10.1113/jphysiol.2005.086280

20. Zhou L, Aon MA, Liu T, O’Rourke B. Dynamic modulation of Ca2+ sparks by mitochondrial oscillations in isolated guinea pig cardiomyocytes under oxidative stress. Journal of Molecular and Cellular Cardiology. 2011 Nov;51(5):632–9. doi:10.1016/j.yjmcc.2011.05.007

21. Yan Y, Liu J, Wei C, Li K, Xie W, Wang Y, et al. Bidirectional regulation of Ca2+ sparks by mitochondria-derived reactive oxygen species in cardiac myocytes. Cardiovascular Research. 2008 Jan 15;77(2):432–41. doi:10.1093/cvr/cvm047

22. Tang TH, Chang CT, Wang HJ, Erickson JD, Reichard RA, Martin AG, et al. Oxidative stress disruption of receptor-mediated calcium signaling mechanisms. J Biomed Sci. 2013 Jul 12;20(1):48. doi:10.1186/1423-0127-20-48 PubMed PMID: 23844974; PubMed Central PMCID: PMC3716919.

23. Yu Z, Wang H, Tang W, Wang S, Tian X, Zhu Y, et al. Mitochondrial Ca2+ oscillation induces mitophagy initiation through the PINK1-Parkin pathway. Cell Death Dis. 2021 Jun 19;12(7):1–7. doi:10.1038/s41419-021-03913-3

24. Baker MR, Fan G, Serysheva II. Structure of IP3R channel: high-resolution insights from cryo-EM. Current Opinion in Structural Biology. 2017 Oct;46:38–47. doi:10.1016/j.sbi.2017.05.014

25. Kang S, Kang J, Kwon H, Frueh D, Yoo SH, Wagner G, et al. Effects of redox potential and Ca2+ on the inositol 1,4,5-trisphosphate receptor L3-1 loop region: implications for receptor regulation. J Biol Chem. 2008 Sep 12;283(37):25567–75. doi:10.1074/jbc.M803321200 PubMed PMID: 18635540.

26. Bánsághi S, Golenár T, Madesh M, Csordás G, RamachandraRao S, Sharma K, et al. Isoform-and Species-specific Control of Inositol 1,4,5-Trisphosphate (IP3) Receptors by Reactive Oxygen Species. Journal of Biological Chemistry. 2014 Mar;289(12):8170–81. doi:10.1074/jbc.M113.504159

27. Lock JT, Sinkins WG, Schilling WP. Protein S-glutathionylation enhances Ca2+-induced Ca2+ release via the IP3 receptor in cultured aortic endothelial cells. J Physiol. 2012 Aug 1;590(15):3431–47. doi:10.1113/jphysiol.2012.232645 PubMed PMID: 22855054; PubMed Central PMCID: PMC3547261.

28. Joseph SK, Young MP, Alzayady K, Yule DI, Ali M, Booth DM, et al. Redox regulation of type-I inositol trisphosphate receptors in intact mammalian cells. J Biol Chem. 2018 Nov 9;293(45):17464–76. doi:10.1074/jbc.RA118.005624 PubMed PMID: 30228182; PubMed Central PMCID: PMC6231128.

29. Ehring GR, Kerschbaum HH, Fanger CM, Eder C, Rauer H, Cahalan MD. Vanadate induces calcium signaling, Ca2+ release-activated Ca2+ channel activation, and gene expression in T lymphocytes and RBL-2H3 mast cells via thiol oxidation. J Immunol. 2000 Jan 15;164(2):679–87. doi:10.4049/jimmunol.164.2.679 PubMed PMID: 10623810.

30. Thrower EC, Duclohier H, Lea EJA, Molle G, Dawson AP. The inositol 1,4,5-trisphosphate-gated Ca2+ channel: effect of the protein thiol reagent thimerosal on channel activity. Biochemical Journal. 1996 Aug 15;318(1):61–6. doi:10.1042/bj3180061

31. Redondo PC, Salido GM, Rosado JA, Pariente JA. Effect of hydrogen peroxide on Ca2+ mobilisation in human platelets through sulphydryl oxidation dependent and independent mechanisms. Biochem Pharmacol. 2004 Feb 1;67(3):491–502. doi:10.1016/j.bcp.2003.09.031 PubMed PMID: 15037201.

32. Höfer T. Model of Intercellular Calcium Oscillations in Hepatocytes: Synchronization of Heterogeneous Cells. Biophysical Journal. 1999 Sep;77(3):1244–56. doi:10.1016/S0006-3495(99)76976-6

33. Catacuzzeno L. A theoretical study on the role of Ca2+-activated K+ channels in the regulation of hormone-induced Ca2+ oscillations and their synchronization in adjacent cells. Journal of Theoretical Biology. 2012;10.

34. Cao P, Donovan G, Falcke M, Sneyd J. A Stochastic Model of Calcium Puffs Based on Single-Channel Data. Biophys J. 2013 Sep 3;105(5):1133–42. doi:10.1016/j.bpj.2013.07.034 PubMed PMID: 24010656; PubMed Central PMCID: PMC3852038.

35. Qi H, Li L, Shuai J. Optimal microdomain crosstalk between endoplasmic reticulum and mitochondria for Ca2+ oscillations. Sci Rep. 2015 Jan 23;5(1):1. doi:10.1038/srep07984

36. Wacquier B, Combettes L, Van Nhieu GT, Dupont G. Interplay Between Intracellular Ca2+ Oscillations and Ca2+-stimulated Mitochondrial Metabolism. Sci Rep. 2016 May 10;6(1):19316. doi:10.1038/srep19316

37. Han JM, Periwal V. A mathematical model of calcium dynamics: Obesity and mitochondria-associated ER membranes. Sneyd J, editor. PLoS Comput Biol. 2019 Aug 22;15(8):e1006661. doi:10.1371/journal.pcbi.1006661

38. Moshkforoush A, Ashenagar B, Tsoukias NM, Alevriadou BR. Modeling the role of endoplasmic reticulum-mitochondria microdomains in calcium dynamics. Sci Rep. 2019 Dec;9(1):17072. doi:10.1038/s41598-019-53440-7

39. Manhas N. A mathematical model of intricate calcium dynamics and modulation of calcium signalling by mitochondria in pancreatic acinar cells. 2021;19.

40. Yang XS. Computational modelling of nonlinear calcium waves. Applied Mathematical Modelling. 2006 Feb 1;30(2):200–8. doi:10.1016/j.apm.2005.03.013

41. Ornelas-Guevara R, Diercks BP, Guse AH, Dupont G. Ca2+ puffs underlie adhesion-triggered Ca2+ microdomains in T cells. Biochimica et Biophysica Acta (BBA) - Molecular Cell Research. 2024 Dec;1871(8):119808. doi:10.1016/j.bbamcr.2024.119808

42. Gil D, Guse AH, Dupont G. Three-Dimensional Model of Sub-Plasmalemmal Ca2+ Microdomains Evoked by the Interplay Between ORAI1 and InsP3 Receptors. Front Immunol. 2021 Apr 28;12:659790. doi:10.3389/fimmu.2021.659790

43. Pages N, Vera-Sigüenza E, Rugis J, Kirk V, Yule DI, Sneyd J. A Model of $$\hbox {Ca}^{2+}$$ Dynamics in an Accurate Reconstruction of Parotid Acinar Cells. Bull Math Biol. 2019 May;81(5):1394–426. doi:10.1007/s11538-018-00563-z

44. Sneyd J, Han JM, Wang L, Chen J, Yang X, Tanimura A, et al. On the dynamical structure of calcium oscillations. Proc Natl Acad Sci USA. 2017 Feb 14;114(7):1456–61. doi:10.1073/pnas.1614613114

45. Li Q, Su D, O’Rourke B, Pogwizd SM, Zhou L. Mitochondria-derived ROS bursts disturb Ca ^2+^ cycling and induce abnormal automaticity in guinea pig cardiomyocytes: a theoretical study. American Journal of Physiology-Heart and Circulatory Physiology. 2015 Mar 15;308(6):H623–36. doi:10.1152/ajpheart.00493.2014

46. Song Z, Xie LH, Weiss JN, Qu Z. A Spatiotemporal Ventricular Myocyte Model Incorporating Mitochondrial Calcium Cycling. Biophysical Journal. 2019 Dec;117(12):2349–60. doi:10.1016/j.bpj.2019.09.005

47. Dupont G, Falcke M, Kirk V, Sneyd J. Models of Calcium Signalling. Vol. 43. Cham: Springer International Publishing; 2016. (Interdisciplinary Applied Mathematics). doi:10.1007/978-3-319-29647-0

48. Watanabe S, Ilieva H, Tamada H, Nomura H, Komine O, Endo F, et al. Mitochondria-associated membrane collapse is a common pathomechanism in SIGMAR1- and SOD1-linked ALS. EMBO Mol Med. 2016 Dec;8(12):1421–37. doi:10.15252/emmm.201606403 PubMed PMID: 27821430; PubMed Central PMCID: PMC5167132.

49. van Stroe-Biezen SAM, Everaerts FM, Janssen LJJ, Tacken RA. Diffusion coefficients of oxygen, hydrogen peroxide and glucose in a hydrogel. Analytica Chimica Acta. 1993 Feb 15;273(1):553–60. doi:10.1016/0003-2670(93)80202-V

50. Johri A, Chandra A. Connection Lost, MAM: Errors in ER–Mitochondria Connections in Neurodegenerative Diseases. Brain Sciences. 2021 Nov;11(11):11. doi:10.3390/brainsci11111437

51. Khan SA, Rossi AM, Riley AM, Potter BVL, Taylor CW. Subtype-selective regulation of IP3 receptors by thimerosal via cysteine residues within the IP3-binding core and suppressor domain. Biochemical Journal. 2013 Mar 28;451(2):177–84. doi:10.1042/BJ20121600

52. Bánsághi S, Golenár T, Madesh M, Csordás G, RamachandraRao S, Sharma K, et al. Isoform- and Species-specific Control of Inositol 1,4,5-Trisphosphate (IP3) Receptors by Reactive Oxygen Species *. Journal of Biological Chemistry. 2014 Mar 14;289(12):8170–81. doi:10.1074/jbc.M113.504159

53. Wüst RCI, Helmes M, Martin JL, van der Wardt TJT, Musters RJP, van der Velden J, et al. Rapid frequency-dependent changes in free mitochondrial calcium concentration in rat cardiac myocytes. The Journal of Physiology. 2017;595(6):2001–19. doi:10.1113/JP273589

54. Okuda M, Tsuruta T, Katayama K. Lifetime and diffusion coefficient of active oxygen species generated in TiO2 sol solutions. Phys Chem Chem Phys. 2009 Mar 18;11(13):2287–92. doi:10.1039/B817695G

55. Li X, Zhang P, Yin Z, Xu F, Yang ZH, Jin J, et al. Caspase-1 and Gasdermin D Afford the Optimal Targets with Distinct Switching Strategies in NLRP1b Inflammasome-Induced Cell Death. Research (Wash D C). 2022 Jul 19;2022:9838341. doi:10.34133/2022/9838341 PubMed PMID: 35958114; PubMed Central PMCID: PMC9343085.

56. Li X, Zhong CQ, Wu R, Xu X, Yang ZH, Cai S, et al. RIP1-dependent linear and nonlinear recruitments of caspase-8 and RIP3 respectively to necrosome specify distinct cell death outcomes. Protein Cell. 2021 Nov;12(11):858–76. doi:10.1007/s13238-020-00810-x PubMed PMID: 33389663; PubMed Central PMCID: PMC8563874.

57. Kirichok Y, Krapivinsky G, Clapham DE. The mitochondrial calcium uniporter is a highly selective ion channel. Nature. 2004 Jan;427(6972):360–4. doi:10.1038/nature02246

58. Boyman L, Lederer WJ. How the mitochondrial calcium uniporter complex (MCUcx) works. Proceedings of the National Academy of Sciences. 2020 Sep 15;117(37):22634–6. doi:10.1073/pnas.2015886117

59. Zhao W bin, Sheng R. The correlation between mitochondria-associated endoplasmic reticulum membranes (MAMs) and Ca2+ transport in the pathogenesis of diseases. Acta Pharmacol Sin. 2025 Feb;46(2):271–91. doi:10.1038/s41401-024-01359-9

60. Csordás G, Renken C, Várnai P, Walter L, Weaver D, Buttle KF, et al. Structural and functional features and significance of the physical linkage between ER and mitochondria. J Cell Biol. 2006 Sep 25;174(7):915–21. doi:10.1083/jcb.200604016 PubMed PMID: 16982799; PubMed Central PMCID: PMC2064383.

61. Arruda AP, Pers BM, Parlakgül G, Güney E, Inouye K, Hotamisligil GS. Chronic enrichment of hepatic endoplasmic reticulum–mitochondria contact leads to mitochondrial dysfunction in obesity. Nat Med. 2014 Dec;20(12):1427–35. doi:10.1038/nm.3735

62. Zandieh A, Shariatpanahi SP, Ravassipour AA, Azadipour J, Nezamtaheri MS, Habibi-Kelishomi Z, et al. An amplification mechanism for weak ELF magnetic fields quantum-bio effects in cancer cells. Sci Rep. 2025 Jan 23;15(1):2964. doi:10.1038/s41598-025-87235-w

63. Zamora-Sillero E, Hafner M, Ibig A, Stelling J, Wagner A. Efficient characterization of high-dimensional parameter spaces for systems biology. BMC Syst Biol. 2011 Sep 15;5(1):142. doi:10.1186/1752-0509-5-142

