## Supplementary Information for "A Dynamic Model of Calcium Frequency Modulation by Mitochondrial Superoxide in Non-excitable Cells"

**For**

### Supplementary Methods

#### 1. Compartmentalization

Our modeling was based on a system of ordinary differential equations (ODEs). The model developed with six compartments: 1) the matrix of a single mitochondrion ($V_M$), 2) the cytosolic region between a mitochondrion and adjacent endoplasmic reticulum (ER) membrane known as mitochondria-associated membrane (MAM) ($V_{MAM}$), which we refer to briefly as the MAM cytosol, 3) the peripheral cytosolic microdomain of a single mitochondrion ($V_N$) excluding $V_{MAM}$, which hereafter we refer to briefly as the near cytosol, 4) the bulk cytosol ($V_B$), 5) the lumen of ER adjacent to a mitochondrion ($V_{EN}$), and 6) the lumen of endoplasmic reticulum within the bulk cytoplasm ($V_{EB}$) (Fig S1).

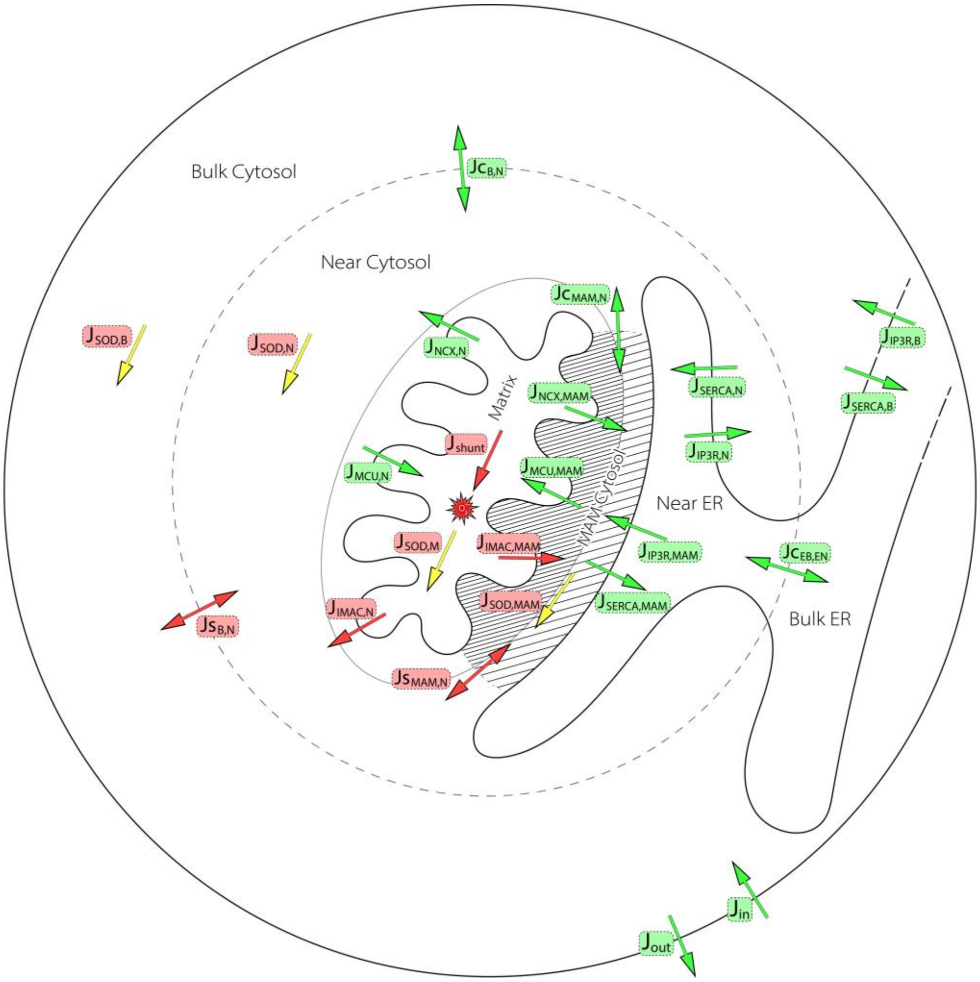


*Fig S1: Fluxes/reactions of superoxide and calcium. Red highlights indicate superoxide-related processes, while green highlights correspond to calcium-related processes.*

Due to the rather high permeability of the outer mitochondrial membrane to ions, the intermembrane space of the mitochondrion is omitted in this model, as in similar models (1,2), and treated as part of the cytosolic space.

As in the previous studies (3,4), we assumed that the endoplasmic reticulum is uniformly distributed throughout the cytoplasm, with a volumetric ratio of 1 to 5.5 (Volume of ER to cytosol). Finally, the bulk cytosolic volume was estimated based on the average volume of spherical cells, taken to be 1000 μm³.

#### 2. Radius and volume of mitochondria and the near cytosol:

We used electron microscopy images (5) to estimate the mitochondrial volume. Accordingly, the mitochondrial matrix approximated as a sphere with a radius of 0.3 μm ($R_{mit}$), and its volume calculated based on this assumption (Fig S2, Eq S3).

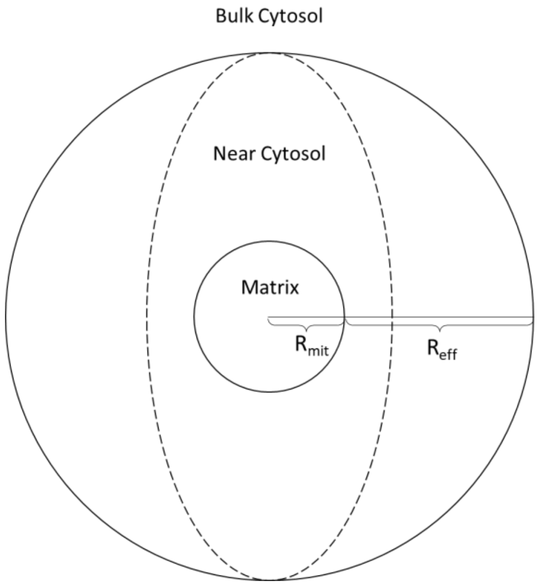


*Fig S2: A single mitochondrion and near cytosol.*

The radius of the near cytosol was determined using fluorescence microscopy images (6) and it was estimated to be 1 μm. According to the random walk model, this radius corresponds to:

$$(S1) \quad R_{eff,S} = \sqrt{D_S \times \tau_S}$$

$\tau_S$ is the half-life of superoxide. Therefore, based on its enzymatic degradation, it can be expressed as given in Eq S2:

$$(S2) \quad \tau_s = \frac{K_{SOD}}{k_{SOD}}$$

The diffusion coefficient of hydrogen peroxide in a gel-like environment has been reported as 370 μm²/s (7). Given the radical and ionic nature of superoxide, its diffusion coefficient was considered to be significantly lower than that of peroxide, and was set to 10 μm²/s. Accordingly, $\tau_S$ was assumed to be 0.1 s. Thus, the volume of the near cytosol is obtained by the volume of cytosolic sphere (Eq S4):

$$(S3) \quad V_M = \frac{4}{3}\pi\,(R_{mit})^3$$

$$(S4) \quad V_N = \frac{4}{3}\pi\,\left(R_{mit} + R_{eff}\right)^3$$

#### 3. Volumetric ratios

The volume ratios between different compartments of the model—including the mitochondrial matrix ($V_M$), MAM-associated cytosol ($V_{MAM}$), near cytosol ($V_N$), bulk cytosol ($V_B$), endoplasmic reticulum in near cytoplasm ($V_{EN}$), endoplasmic reticulum in bulk cytoplasm ($V_{EB}$), and the extracellular space ($V_{EX}$)—were defined as follows:

$$(S5) \quad \alpha = \frac{V_N}{V_B} = \frac{V_{EN}}{V_{EB}}$$

$$(S6) \quad \beta = \frac{V_B}{V_{EB}} = \frac{V_N}{V_{EN}}$$

$$(S7) \quad \gamma = \frac{V_N}{V_M}$$

$$(S8) \quad \delta = \frac{V_N}{V_{MAM}}$$

$$(S9) \quad \varepsilon = \frac{V_{EX}}{V_B}$$

#### 4. Superoxide dynamics

Given that phospholipid membranes are impermeable to ions, the dynamics of superoxide concentration considered only within the mitochondrial matrix ($S_M$), the MAM cytosol ($S_{MAM}$), the near cytosol ($S_N$), and the bulk cytosol ($S_B$). Furthermore, due to the minimal effect of calcium flux into or out of the matrix on the mitochondrial membrane potential ($\Delta\psi$) (8), we considered only the impact of the opening of the mitochondrial inner membrane anion channels (IMACs) on the mitochondrial membrane potential, following the approach of Yang et al. (9).

$$(S10)\quad \frac{dS_M}{dt} = J_{shunt} - J_{SOD_M} - \gamma.(1 - A_{Mit}).J_{IMAC_N} - \frac{\gamma}{\delta}.A_{Mit}.J_{IMAC_{MAM}}$$

$$(S11)\quad \frac{dS_{MAM}}{dt} = A_{Mit}.J_{IMAC_{MAM}} - J_{SODc_{MAM}} - \delta.J_{S_{MAM,N}}$$

$$(S12)\quad \frac{dS_N}{dt} = (1 - A_{Mit}).J_{IMAC_N} - J_{SODc_N} + J_{S_{MAM,N}} + J_{S_{B,N}}$$

$$(S13)\quad \frac{dS_B}{dt} = -J_{SODc_B} - \alpha.J_{S_{B,N}}$$

And for the mitochondrial membrane potential:

$$(S14)\quad \frac{d\Delta\psi}{dt} = J_{\psi_S} - J_{\psi_U} - (1 - A_{Mit}).J_{\psi_{IMAC_N}} - A_{Mit}.J_{\psi_{IMAC_{MAM}}}$$

The coefficients $A_{Mit}$ and $A_{ER}$ ($A_{ER}$ will be used in the calcium equations) represent the ratios of the membrane surface areas of a mitochondrion and the endoplasmic reticulum that are associated with the MAM cytosol to the total surface area of each organelle membrane (Eq 8 and 9):

$$(S15)\quad A_{ER} = \frac{MAM\ Surface\ Area}{Total\ ER\ Surface\ Area}$$

$$(S16)\quad A_{Mit} = \frac{MAM\ Related\ Mitochondrion\ Surface\ Area}{Total\ Mitochondrion\ Surface\ Area}$$

#### 5. Superoxide Fluxes/Reactions

In the model developed by Yang et al. (9), peroxide dynamics also influence superoxide dynamics through its effect on the mitochondrial permeability transition pores (mPTP). However, to simplify the superoxide model—and given that the activation of mPTP channels predominantly occurs under pathological conditions—peroxide dynamics were omitted in the present study. Consequently, superoxide production in the mitochondrial matrix ($J_{shunt}$) was assumed to occur at a constant rate defined as $k_{shunt}$ (9):

$$(S17)\quad J_{shunt} = k_{shunt}$$

The algebraic definitions of the cytosolic superoxide dismutation rate ($J_{SOD,C}$), mitochondrial superoxide dismutation rate ($J_{SOD,M}$), superoxide leakage through IMAC channels ($J_{IMAC}$), the rate of mitochondrial membrane potential production by cellular respiration ($J_{\psi,S}$), baseline membrane potential leakage rate ($J_{\psi,U}$), and membrane potential leakage due to IMACs opening ($J_{\psi,IMAC}$) were all adopted from the model of Yang et al. (9) and are as follows:

Superoxide flux through the IMAC channel (9):

$$(S18)\quad J_{IMAC_x} = k_{IMAC}\ P_{IMAC_x}\ S_M$$

In this equation, $S_M$ denotes the concentration of superoxide in the mitochondrial matrix, and $P_{IMAC}$ represents the open probability of IMAC channels. Since the open probability of IMAC channels is a function of cytosolic superoxide concentration, the subscript x is used to distinguish between IMAC channels located on the MAM-facing side of the mitochondrial membrane and those on the side facing the near cytosol.

Open probability of IMAC channels (9):

$$(S19)\quad P_{IMAC_x} = 0.001 + 0.999\ \frac{S_x^{h_{IMAC}}}{S_x^{h_{IMAC}} + K_{S_{IMAC}}^{h_{IMAC}}}$$

Enzymatic dismutation of superoxide in the mitochondrial matrix (9):

$$(S20)\quad J_{SOD_m} = k_{SOD_m}\ \frac{S_M}{S_M + K_{SOD_m}}$$

Superoxide dismutation in the cytosol (including all three cytosolic compartments: MAM cytosol, near cytosol, and bulk cytosol) (9):

$$(S21)\quad J_{SOD_{c_x}} = k_{SOD_c}\ \frac{S_x}{S_x + K_{SOD_c}}$$

Superoxide diffusion between the MAM cytosol and the near cytosol (9):

$$(S22)\quad J_{S_{MAM,N}} = k_{S_{MAM,N}}\ (S_{MAM} - S_N)$$

Superoxide diffusion between the near cytosol and the bulk cytosol (9):

$$(S23)\quad J_{S_{B,N}} = k_{S_{B,N}}\ (S_B - S_N)$$

Potential dependent leakage of mitochondrial membrane potential (9):

$$(S24)\quad J_{\psi_U} = k_{\psi_U}\ \Delta\psi$$

Membrane potential leakage due to IMAC channel opening (9):

$$(S25) \quad J_{\psi_{IMAC_x}} = k_{\psi_{IMAC}} P_{IMAC_x} \Delta\psi$$

The subscript x indicates that this equation is written separately for IMAC channels located on the MAM-associated side and the other regions of the mitochondrial membrane.

#### 6. Diffusion constants of superoxide and calcium between cytosolic compartments

To calculate the diffusion constants of superoxide and calcium between the MAM cytosol and the near cytosol, denoted as $k_{S,MAM,N}$ and $k_{C,MAM,N}$, respectively, we employed estimations based on Fick's law. The MAM cytosol was first approximated as a rectangular cuboid, in which the width of the cuboid corresponds to the distance between a mitochondrion and the endoplasmic reticulum, maintained by specific tethering proteins (Fig S3). Based on microscopic observations (5), this distance was determined to be 20 nm.

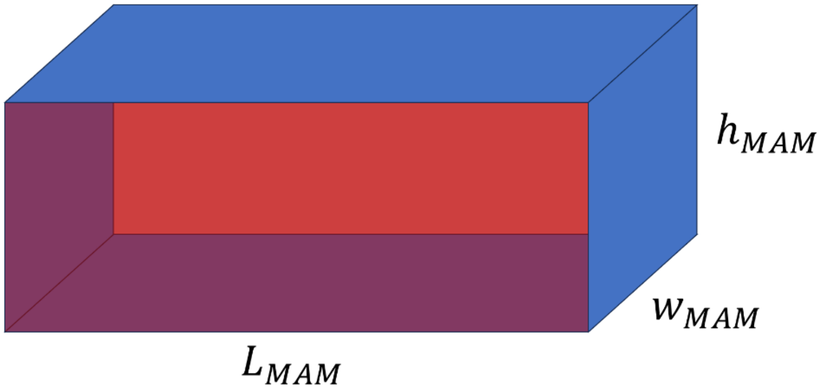


*Fig S3: Approximation of the MAM cytosol as a rectangular cuboid. The blue faces represent the surfaces through which free diffusion of species occurs between the MAM cytosol and the near cytosol. One of the red faces corresponds to the mitochondrial membrane, while the opposite red face represents the MAM region of ER.*

The length ($L_{MAM}$) of the MAM cytosol was set to 200 nm based on experimental observations (5), and its height ($h_{MAM}$) was set as half that value, i.e., 100 nm.
Using integrated form of Fick's law, the diffusion constants of superoxide and calcium between the MAM cytosol and the near cytosol can now be estimated by Eq S26 and S27, respectively:

$$(S26) \quad k_{S_{MAM,N}} = D_S \frac{A_{MAM}}{l_{MAM}} \frac{1}{V_N}$$

$$(S27) \quad k_{C_{MAM,N}} = D_{Ca} \frac{A_{MAM}}{l_{MAM}} \frac{1}{V_N}$$

In these equations, $D_S$ and $D_{Ca}$ represent the diffusion coefficients of superoxide and calcium in the cytosol, respectively. $A_{MAM}$ denotes the total surface area through which the near cytosol and the MAM cytosol are connected (the blue surfaces in Fig S3). Note that the $l_{MAM}$ in Eq S26 and S27 is a different parameter from previously defined $L_{MAM}$ and is the distance from the center of the MAM space to its hypothetical boundaries, taken as the smaller value between $h_{MAM}$ and $L_{MAM}$.
To calculate the diffusion constant of superoxide between the near cytosol and the bulk cytosol ($k_{S,B,N}$), Fick's law was applied considering the geometry of the mitochondrion and the near cytosol (Fig S2), using Eq S28:

$$(S28) \quad k_{S_{B,N}} = 4\pi\, D_S \left( \frac{1}{\frac{1}{R_{mit}} - \frac{1}{R_{eff}}} \right) \frac{1}{V_N}$$

To calculate the corresponding diffusion constant for calcium, considering the tubular structure of the ER (Fig S4) and based on the Fick's law, Eq S29 was used to estimate the constant $k_{C,B,N}$. The radius of the cross-sectional area of the endoplasmic reticulum (denoted as a in Fig S4) was determined to be 20 nm based on electron microscopy images (5). The value of b was also considered to be approximately equal to L/2 in Fig S3:

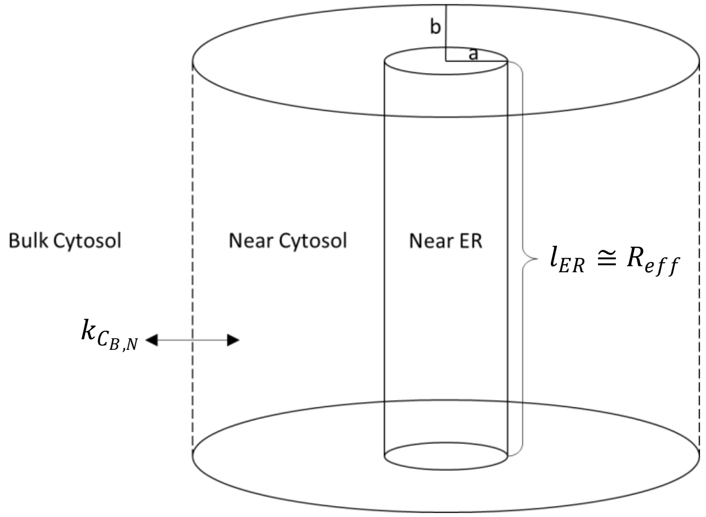


*Fig S4: Approximation of the endoplasmic reticulum as a cylinder for estimating the diffusion constant of calcium between the near cytosol and bulk cytosol. The dashed lines indicate the sites of calcium diffusion.*

$$(S29)\quad k_{C_{B,N}} = 2\pi\, D_{Ca}\, \frac{L}{\ln\left(\frac{b}{a}\right)} \frac{1}{V_N} = 2\pi\, D_{Ca}\, \frac{2\, R_{eff}}{\ln\left(\frac{R_{eff}}{a}\right)} \frac{1}{V_N}$$

Finally, to calculate the diffusion constant of calcium within the lumen of ER present in the near cytoplasm and the bulk cytoplasm ($k_{C,EB,E}$), the same cylindrical geometry was applied to both compartments (Fig S5). Eq S30 provides the estimation of this diffusion constant:

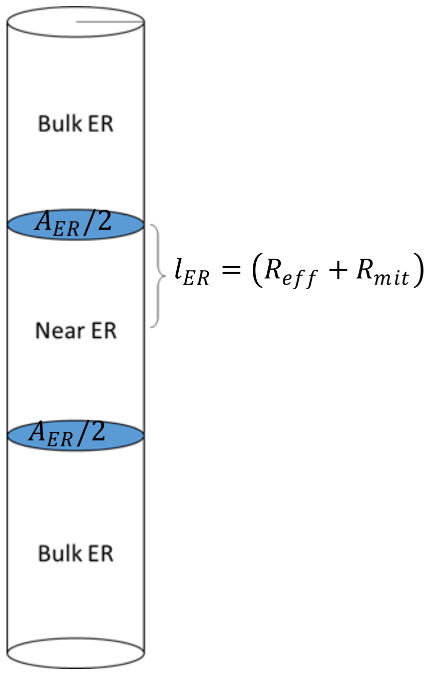


*Figure S5: Approximation of the near and bulk endoplasmic reticulum as cylinders. The blue discs represent the hypothetical boundaries across which calcium diffuses between these two compartments.*

$$(S30)\quad k_{Ce_{B,N}} = D_{ca} \frac{A_{ER}}{l_{ER}} \frac{1}{V_{ER_N}} = D_{Ca} \frac{2\,\pi\, a^2}{(R_{eff} + R_{mit})} \frac{\beta}{V_{CN}}$$

#### 7. Calcium Dynamics

We used open-cell condition and Class 1 dynamics of calcium in our model that means calcium oscillations can occur at constant concentrations of inositol 1,4,5-trisphosphate (IP3) as the primary stimulant of inositol 1,4,5-trisphosphate receptors (IP3R) (4). Furthermore, consistent with previous models (1,4), the rapid buffering approximation are adopted for calcium in both the cytosol and the mitochondrial matrix (represented by the coefficient $f_m$ in Eq S38), and thus omitted the explicit dynamics of calcium buffering.

IP3R channels on the endoplasmic reticulum (ER) membrane accumulate at the MAM region through protein linkages with voltage-dependent anion channels (VDACs) located on the outer mitochondrial membrane (10). This leads to a non-uniform calcium flux into the mitochondrial matrix via mitochondrial calcium uniporters (MCUs), which reside in the inner mitochondrial membrane. As a result, the calcium flux per unit mitochondrial membrane area is elevated in the MAM region. Therefore, using only $A_{ER}$ (for IP3Rs) and $A_{Mit}$ (for MCUs) is insufficient to account for the non-uniform distribution of these channels across membrane regions. Thus, the additional coefficients are defined as follows:

$$(S31) \quad W_{IP3R} = \frac{Density\ of\ IP3R\ at\ MAM}{Density\ of\ IP3R\ at\ Other\ Reigons\ of\ ER\ Membrane}$$

$$(S32) \quad W_{MCU} = \frac{Density\ of\ MCU\ at\ MAM\ Related\ Mitochondrial\ Membrane}{Density\ of\ MCU\ at\ Other\ Reigons\ of\ Mitochondrial\ Membrane}$$

The equations governing the free calcium dynamics in the bulk cytosol ($C_B$), the endoplasmic reticulum within the bulk cytoplasm ($C_{EB}$), the near cytosol ($C_N$), the endoplasmic reticulum adjacent to a mitochondrion ($C_{EN}$), the cytosol within the MAM region ($C_{MAM}$), and the mitochondrial matrix ($C_M$) are as follows:

$$(S33) \quad \frac{dC_B}{dt} = (1 - A_{ER})\,.J_{IP3R_B} - (1 - A_{ER})\,.J_{SERCA_B} - \alpha\,.J_{C_{B,N}} + \varepsilon\,.(J_{in} - J_{out})$$

$$(S34) \quad \frac{dCe_B}{dt} = \beta\,.\left[(1 - A_{ER})\,.J_{SERCA_B} - (1 - A_{ER})\,.J_{IP3R_B}\right] - \alpha\,.J_{C_{EB,EN}}$$

$$(S35) \quad \frac{dC_N}{dt} = (1 - A_{ER})\,.J_{IP3R_N} - (1 - A_{ER})\,.J_{SERCA_N}$$

$$+ \frac{1}{\gamma}\,.\left[(1 - A_{Mit})\,.J_{NCX_N} - (1 - A_{Mit})\,.J_{MCU_N}\right] + J_{C_{MAM,N}} + J_{C_{B,N}}$$

$$(S36) \quad \frac{dCe_N}{dt} = \beta\,.\left[(1 - A_{ER})\,.J_{SERCA_N} - (1 - A_{ER})\,.J_{IP3R_N}\right]$$

$$+ \frac{\beta}{\delta}\,.\left[A_{ER}\,.J_{SERCA_{MAM}} - A_{ER}.W_{IP3R}\,.J_{IP3R_{MAM}}\right] + J_{C_{EB,EN}}$$

$$(S37) \quad \frac{dC_{MAM}}{dt} = A_{ER}.W_{IP3R}\,.J_{IP3R_{MAM}} - A_{ER}\,.J_{SERCA_{MAM}}$$

$$+ \frac{\delta}{\gamma}\,.\left[A_{Mit}\,.J_{NCX_{MAM}} - A_{Mit}.W_{MCU}\,.J_{MCU_{MAM}}\right] - \delta\,.J_{C_{MAM,N}}$$

$$(S38) \quad \frac{dC_M}{dt} = f_m\,.\left[(1 - A_{Mit})\,.J_{MCU_N} - (1 - A_{Mit})\,.J_{NCX_N} + A_{Mit}.W_{MCU}\,.J_{MCU_{MAM}}\right.$$

$$\left. - A_{Mit}\,.J_{NCX_{MAM}}\right]$$

The equations governing the time-dependent variable h for open probability of IP3R channels located at the MAM region, ER membrane adjacent to mitochondria, and at the ER membrane in bulk cytoplasm are as follows (4):

$$(S39)\quad \frac{dh_{MAM}}{dt} = \frac{h_{\infty_{MAM}} - h_{MAM}}{\tau_{h_{MAM}}}$$

$$(S40)\quad \frac{dh_N}{dt} = \frac{h_{\infty_N} - h_N}{\tau_{h_N}}$$

$$(S41)\quad \frac{dh_B}{dt} = \frac{h_{\infty_B} - h_B}{\tau_{h_B}}$$

#### 8. Calcium Fluxes

Membrane potential-dependent calcium influx into the mitochondrial matrix via mitochondrial calcium uniporter (MCU) (2):

$$(S42)\quad J_{MCU_x} = V_{MCU} \frac{C_x^2}{K_{MCU}^2 + C_x^2}\, \Delta\psi_{MCU}$$

Potential-dependency of MCU flux (11):

$$(S43)\quad \Delta\psi_{MCU} = \exp(P_1\, \Delta\psi)$$

Membrane potential-dependent calcium efflux from the mitochondrial matrix to the surrounding cytosol via mitochondrial sodium-calcium exchanger (NCX) (2):

$$(S44)\quad J_{NCX_x} = V_{NCX} \frac{C_M}{K_{NCX} + C_M} \Delta\psi_{NCX}$$

Potential-dependency of NCX flux (11):

$$(S45)\quad \Delta\psi_{NCX} = \exp(P_2\, \Delta\psi)$$

Calcium diffusion between the MAM cytosol and near cytosol:

$$(S46)\quad J_{C_{MAM,N}} = k_{C_{MAM,N}}\,(C_{MAM} - C_N)$$

Calcium diffusion between the near cytosol and bulk cytosol:

$$(S47)\quad J_{C_{B,N}} = k_{C_{B,N}}\,(C_B - C_N)$$

Calcium diffusion within the lumen of endoplasmic reticulum between the near ER and bulk ER:

$$(S48) \quad J_{C_{EB,EN}} = k_{Ce_{B,N}} \, (Ce_B - Ce_N)$$

Store-operated calcium entry (SOCE) from the plasma membrane (4):

$$(S49) \quad J_{in} = a_0 + a_1 \, \frac{K_e^4}{K_e^4 + C_B^4}$$

Calcium extrusion from the plasma membrane via calcium ATPase pumps (4):

$$(S50) \quad J_{out} = V_{out} \, \frac{C_B^2}{K_{out}^2 + C_B^2}$$

Calcium uptake from cytosol into the ER via the sarco/endoplasmic reticulum calcium ATPase (SERCA) (4):

$$(S51) \quad J_{SERCA_x} = V_{SERCA} \, \frac{(C_x^2 - \bar{K} \, C_{ex}^2)}{C_x^2 + K_{serca}^2}$$

The subscript x denotes SERCA pumps located in the ER of the MAM region, near cytoplasm, and bulk cytoplasm.

Calcium release from ER to the cytosol via IP3R channels (4):

$$(S52) \quad J_{IP3R_x} = k_f \, P_{o_x} \, (C_{ex} - C_x)$$

$P_o$ is the open probability of IP3R channel. The subscript x refers to different regions of ER.

Equations for the IP3R open probability based on Sneyd et al. (4):

$$(S53) \quad P_{o_x} = \frac{\beta_p}{\beta_p + k_\beta \, (\beta_p + \alpha_p)} \quad , \quad (S54) \quad \alpha_p = A \left(1 - \bar{m}_a \, \bar{h}_\alpha\right) \quad , \quad (S55) \quad \beta_p = B \, \bar{m}_\beta \, h \quad ,$$

$$(S56) \quad \bar{m}_a = \bar{m}_\beta = \frac{C_x^4}{K_C^4 + C_x^4} \quad , \quad (S57) \quad \bar{h}_\alpha = h_\infty = \frac{K_h^4}{K_h^4 + C_x^4} \quad , \quad (S58) \quad 1 - A = B = \frac{IP3^2}{K_{P_x}^2 + IP3^2} \quad ,$$

$$(S59) \quad \tau_h = \tau_{max} \, \frac{K_\tau^4}{K_\tau^4 + C_x^4}$$

Here, x represents the different regions of ER and cytosolic compartments and corresponding calcium concentrations.

#### 9. The Proposed Model for The Redox Modulation of IP3Rs Open Probability

In this model, we proposed that superoxide-induced oxidation of IP3R channels directly reduces the half-maximal concentration of IP3 for feedback on IP3Rs, namely $K_P$ (that used in Eq S58), thereby increasing open probability of IP3Rs. To quantify this reduction, we defined a variable called the oxidation fraction of the IP3R channel, as follows:

$$(S60)\quad IP3R_{OX} = \frac{\#\,Oxidized\ Cys\ Residues\ of\ IP3R}{\#\,Total\ Cys\ Resdues\ of\ IP3R}$$

$$(S61)\quad IP3R_{OX} + IP3R_{RED} = 1$$

We then expressed $K_P$ as a function of this relative oxidation fraction using the following equation:

$$(S62)\quad K_P = K_{P_{RED}} \cdot \left(1 - g_{max} \cdot \frac{IP3R_{OX}}{G_{OX} + IP3R_{OX}}\right)$$

Here:

- $K_{P,RED}$ is the value of $K_P$ when the channel is fully reduced (non-oxidized),
- $g_{max}$ denotes the maximum reduction in $K_p$ under full oxidation of channel,
- $G_{OX}$ is the oxidation fraction at which $K_P$ is reduced by half of $g_{max}$.

Fig S6 plots the relationship between $K_P$ and the oxidized fraction of the channels ($IP3R_{OX}$) according to Eq S62:

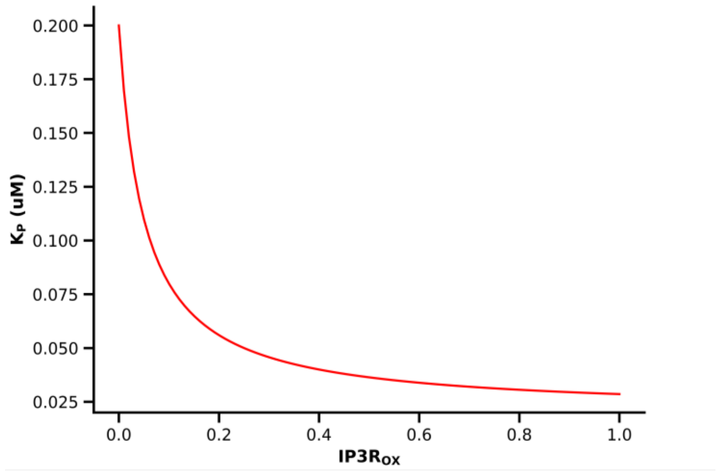


*Fig S6: Illustration of the relationship between $IP3R_{OX}$ and $K_P$*

The premise of the preceding equation, rooted in a quasi-linear assumption, dictates that the $K_p$ value undergoes non-cooperative modifications in response to shifts in the IP3R redox state. However, had this modulation been modeled as a cooperative process, the functional form of $K_p$ would change from a hyperbolic profile to a sigmoidal one, according to Eq S63:

$$(S63) \quad K_P = K_{P_{RED}} \cdot \left(1 - g_{max} \cdot \frac{IP3R_{OX}^{n}}{G_{OX}^{n} + IP3R_{OX}^{n}}\right)$$

$$n = Hill\ coefficient\ \neq 1$$

The distinction between these two models is illustrated in Fig S7.

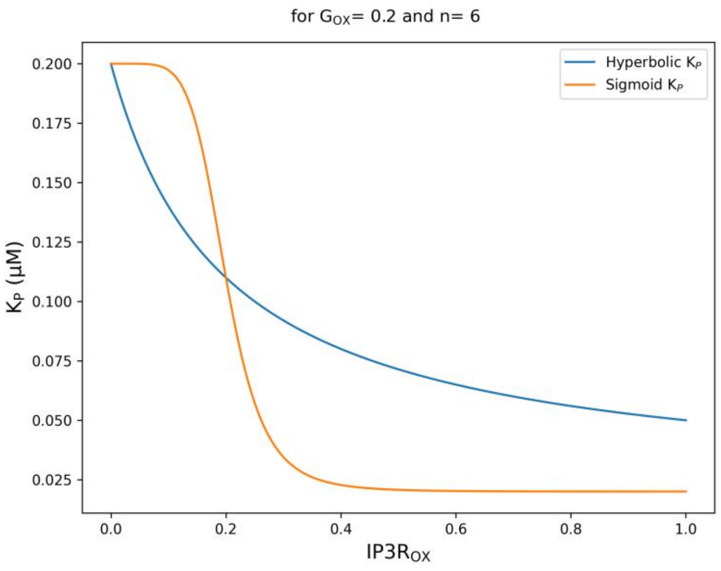


*Fig. S7: Comparison of hyperbolic and sigmoidal functions for modeling $K_p$ modulation.*

In this study, to model the dynamics of the redox state of the IP3R channel, we employed a simple kinetic model. We assumed that the reversible oxidation of the IP3R channel by superoxide follows a mass-action model, and proposed Eq S63 to describe the rate of this oxidation. Additionally, we assumed that the reduction of these channels is enzymatic, mediated by a reducing agent whose concentration remains constant during the superoxide dynamics in the current model; accordingly, we proposed Eq S64 to describe the enzymatic reduction rate of the channel:

$$(S64) \quad V_{OX_x} = k_{OX} \, . S_x \, . IP3R_{RED_x}$$

$$(S65) \quad V_{RED_x} = k_{RED} \cdot \frac{IP3R_{OX_x}}{K_{RED} + IP3R_{OX_x}}$$

Ultimately, the dynamics of the IP3R redox state are described as:

$$(S66)\quad \frac{dIP3R_{OX_{MAM}}}{dt} = V_{OX_{MAM}} - V_{RED_{MAM}}$$

$$(S67)\quad \frac{dIP3R_{OX_N}}{dt} = V_{OX_N} - V_{RED_N}$$

$$(S68)\quad \frac{dIP3R_{OX_B}}{dt} = V_{OX_B} - V_{RED_B}$$

#### 10. Model parameters

Table S1: Model parameters.

| Symbol | Description | Value [unit] | Reference |
|---|---|---|---|
| **Geometrical Parameters** | | | |
| $\boldsymbol{w_{MAM}}$ | MAM cytosol width | 0.02 [µm] | * |
| $\boldsymbol{h_{MAM}}$ | MAM cytosol height | 0.1 [µm] | * |
| $\boldsymbol{L_{MAM}}$ | MAM cytosol length | 0.2 [µm] | * |
| $\boldsymbol{R_{mit}}$ | A single mitochondrion radius | 0.3 [µm] | * |
| $\boldsymbol{V_B}$ | Bulk cytosol volume | 1000 [µm$^3$] | * |
| $\boldsymbol{\beta}$ | ER to cytosol volume ratio | 5.5 | (4) |
| $\boldsymbol{\varepsilon}$ | Extracellular to cellular volume ratio | 1.5 | (4) |
| $\boldsymbol{a}$ | Single ER tube cross-section radius | 0.02 [µm] | * |
| **Superoxide Flux/Reaction Parameters** | | | |
| $\boldsymbol{K_{SOD_C}}$ | Michaelis-Menten constant of cytosolic dismutase | 0.001 [mM] | * |
| $\boldsymbol{k_{SOD_C}}$ | Maximum speed of cytosolic dismutase | 0.01 [mM/s] | * |
| $\boldsymbol{K_{SOD_M}}$ | Michaelis-Menten constant of mitochondrial dismutase | 0.001 [mM] | * |
| $\boldsymbol{k_{SOD_M}}$ | Maximum speed of mitochondrial dismutase | 0.005 [mM/s] | * |
| $\boldsymbol{D_S}$ | Cytosolic diffusion coefficient of superoxide | 10 [µm$^2$/s] | * |
| $\boldsymbol{k_{shunt}}$ | Superoxide production rate in mitochondria matrix | # [mM/s] | * |
| $\boldsymbol{k_{IMAC}}$ | Maximum rate of superoxide flux through IMAC | 0.5 [mM/s] | (9) |
| $\boldsymbol{h_{IMAC}}$ | Hill-coefficient of IMAC for activation by superoxide | 3 | (9) |
| $\boldsymbol{K_{S,IMAC}}$ | Hill-constant of IMAC for activation by superoxide | 0.004 [mM] | (9) |
| $\boldsymbol{k_{\psi_S}}$ | Mitochondrial membrane potential production rate constant | 3.1 [V/s] | * |
| $\boldsymbol{k_{\psi_U}}$ | Mitochondrial membrane (MM) potential leakage rate constant | 19.2 [1/s] | (9) |
| $\boldsymbol{k_{\psi_{IMAC}}}$ | Rate constant of MM potential leakage due to IMAC opening | 5.5 [1/s] | * |

| Calcium Flux Parameters | | | |
|---|---|---|---|
| $\boldsymbol{IP3}$ | IP3 concentration in cytosol | # [μM] | * |
| $\boldsymbol{W_{IP3R}}$ | Density ratio of IP3Rs in MAM to other regions of ER | 7e3 | * |
| $\boldsymbol{\tau_{max}}$ | Time scaling for negative feedback of calcium on IP3R | 35 [1/s] | (4) |
| $\boldsymbol{K_{P_{RED}}}$ | Half-maximal concentration of IP3 for feedback on IP3Rs in full-reduced IP3Rs | 0.2 [μM] | (4) |
| $\boldsymbol{D_{Ca}}$ | Calcium diffusion coefficient in cytosol | 5 [μm$^2$/s] | (3) |
| $\boldsymbol{D_{Ca_e}}$ | Calcium diffusion coefficient in endoplasmic reticulum | 55 [μm$^2$/s] | * |
| $\boldsymbol{k_f}$ | Maximum rate of calcium flux through IP3R | 10 [1/s] | (4) |
| $\boldsymbol{V_{SERCA}}$ | Maximum speed of SERCA | 1.87 [μM/s] | * |
| $\boldsymbol{K_{SERCA}}$ | - | 0.2 | (4) |
| $\boldsymbol{\overline{K}}$ | - | 0.000019 | (4) |
| $\boldsymbol{k_B}$ | - | 0.4 | (4) |
| $\boldsymbol{K_C}$ | Half maximal concentration of calcium for positive feedback on IPR | 0.2 [μM] | (4) |
| $\boldsymbol{K_h}$ | Half maximal concentration of calcium for negative feedback on IPR | 0.08 [μM] | (4) |
| $\boldsymbol{K_\tau}$ | Half maximal concentration of calcium for feedback on time scaling factor | 0.1 [μM] | (4) |
| $\boldsymbol{f_m}$ | Fraction of free calcium in mitochondria matrix | 0.005 | * |
| $\boldsymbol{V_{MCU}}$ | Potential independent maximum speed of MCU | 0.0056 [nM/s] | * |
| $\boldsymbol{K_{MCU}}$ | Hill-constant of MCU activation by calcium | 50 [μM] | * |
| $\boldsymbol{P_1}$ | Constant of MCU potential dependency | 0.1 [1/mV] | (11) |
| $\boldsymbol{W_{MCU}}$ | Density ratio of MCUs in MAM-faced to other regions of mitochondria membrane | 1e5 | * |
| $\boldsymbol{V_{NCX}}$ | Maximum speed of NCX | 6.4 [μM/s] | * |
| $\boldsymbol{K_{NCX}}$ | Half-maximal concentration of calcium for activation of NCX | 30 [μM /s] | * |
| $\boldsymbol{P_2}$ | Constant of NCX potential dependency | 0.016 [1/mV] | (11) |
| $\boldsymbol{V_{OUT}}$ | Maximum speed of calcium efflux through plasma membrane | 0.286 [μM /s] | * |
| $\boldsymbol{K_{OUT}}$ | Half-maximal concentration of calcium for activation of PMCA | 0.3 [μM] | (4) |
| $\boldsymbol{a_0}$ | Constant rate of calcium leakage into cytosol from plasma membrane | 0.0027 [μM /s] | (4) |
| $\boldsymbol{a_1}$ | Maximum flux rate of calcium through SOCE | 0.07 [μM/s] | (4) |
| $\boldsymbol{K_e}$ | Sensitivity of SOCE to bulk ER calcium | 8 | (4) |
| **IP3R Redox Modulation Parameters** | | | |
| $\boldsymbol{k_{OX}}$ | zero-order rate constant for oxidation of IP3Rs by superoxide | 45 [1/s] | * |

| | | | |
|---|---|---|---|
| $\boldsymbol{k_{RED}}$ | maximum reduction rate of oxidized IP3Rs | 0.024 | * |
| $\boldsymbol{K_{RED}}$ | Fraction of oxidized IP3Rs for half-maximum speed of reduction | 0.05 | * |
| $\boldsymbol{g_{max}}$ | maximum decrease in $K_P$ resulting from complete oxidation of IP3Rs | 0.9 | * |
| $\boldsymbol{G_{OX}}$ | oxidized fraction of IP3Rs at which $K_P$ is decreased by half of $g_{max}$ | 0.05 | * |

*: This study.

#: Variable (described in the article).

### Supplementary Results

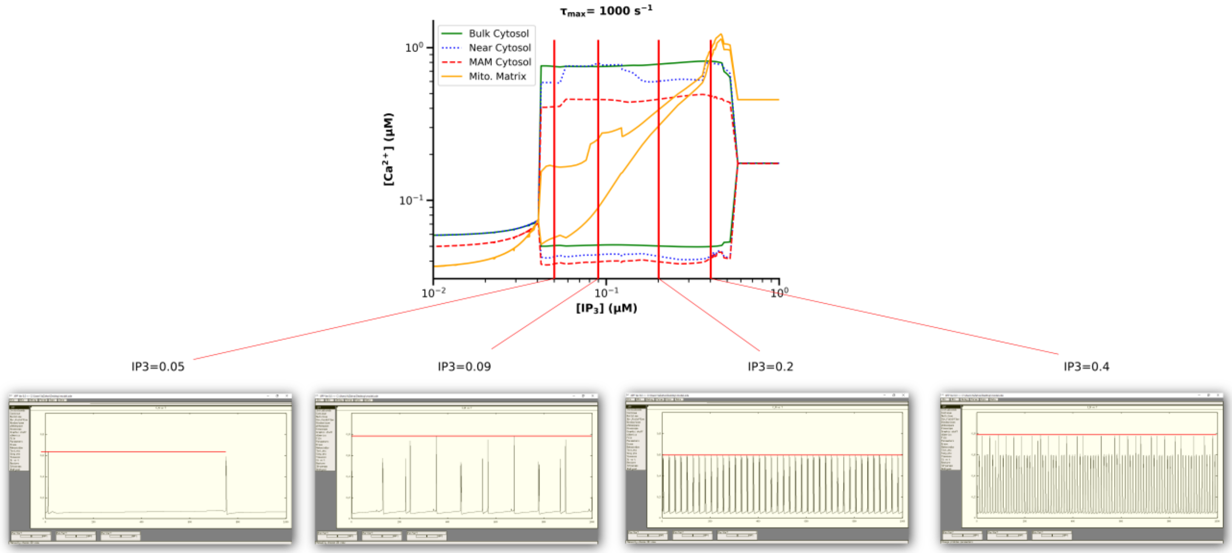


Fig S8: Validation figure for calcium bifurcation (only Near cytosol shown). Four plots in below are the Near cytosol calcium time-series in different concentrations of IP3. Calcium time-series here obtained using XPPAUT.

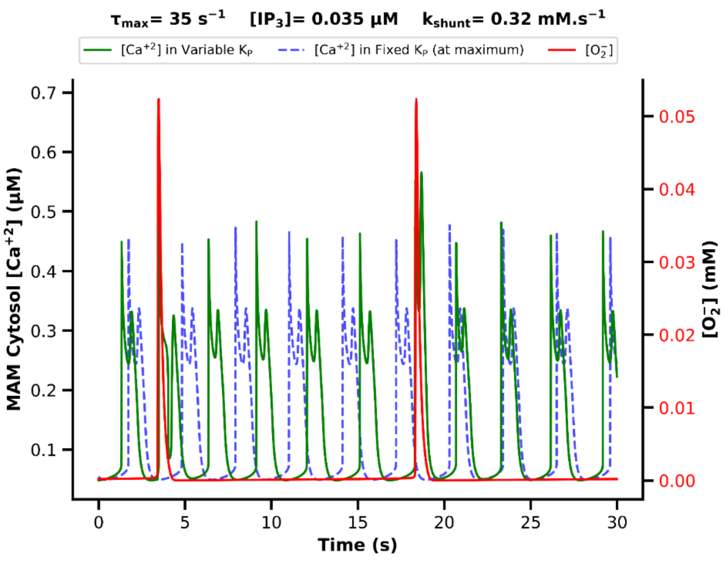


Fig S9: Extended pulse width of calcium oscillation in the MAM cytosol under oscillatory $K_p$ (green curve) versus fixed $K_P$ (blue dashed curve).

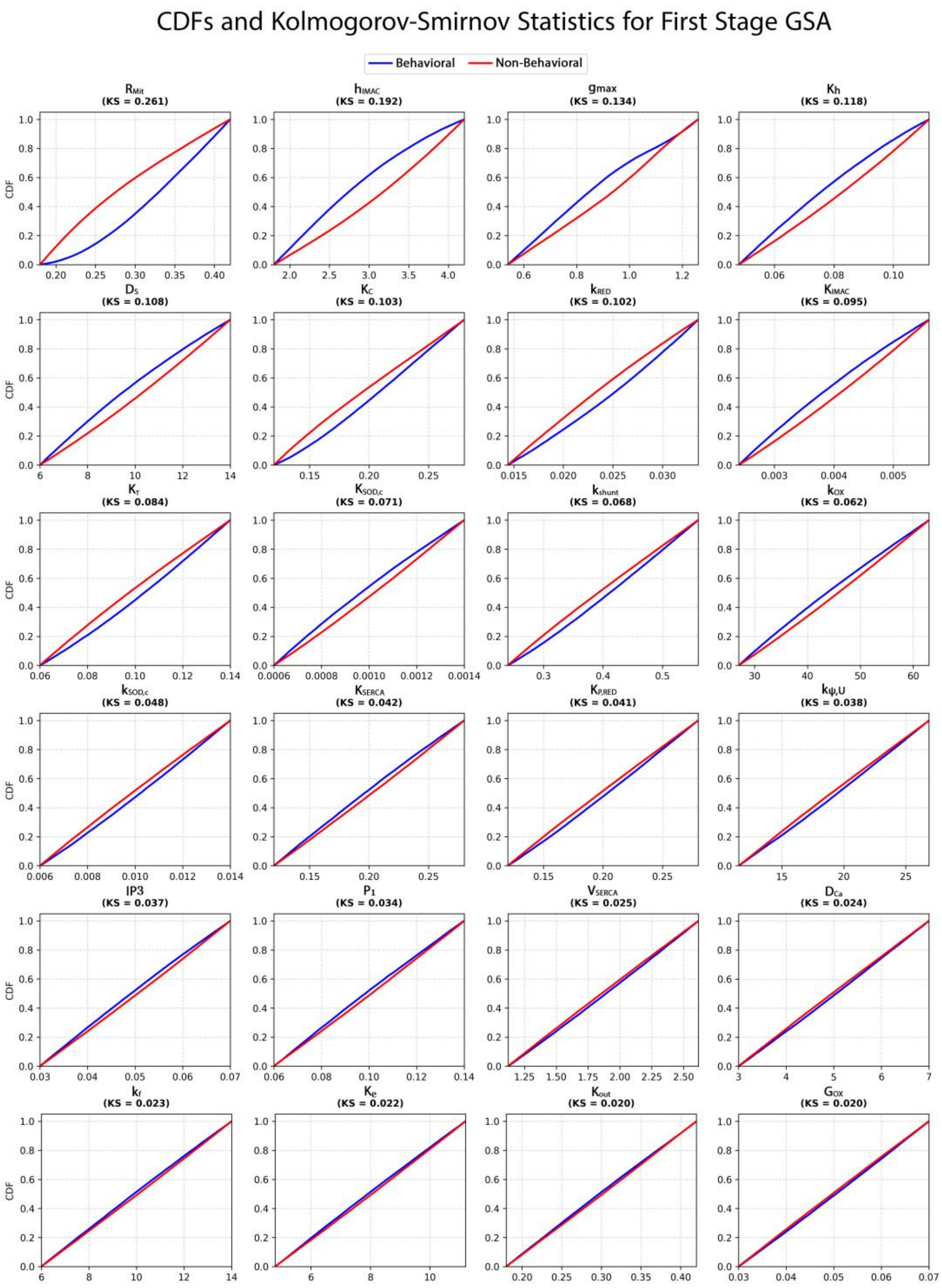


*Fig S10: Cumulative distribution functions plots and Kolmogorov–Smirnov statistics for first-stage global sensitivity analysis of model parameters.*

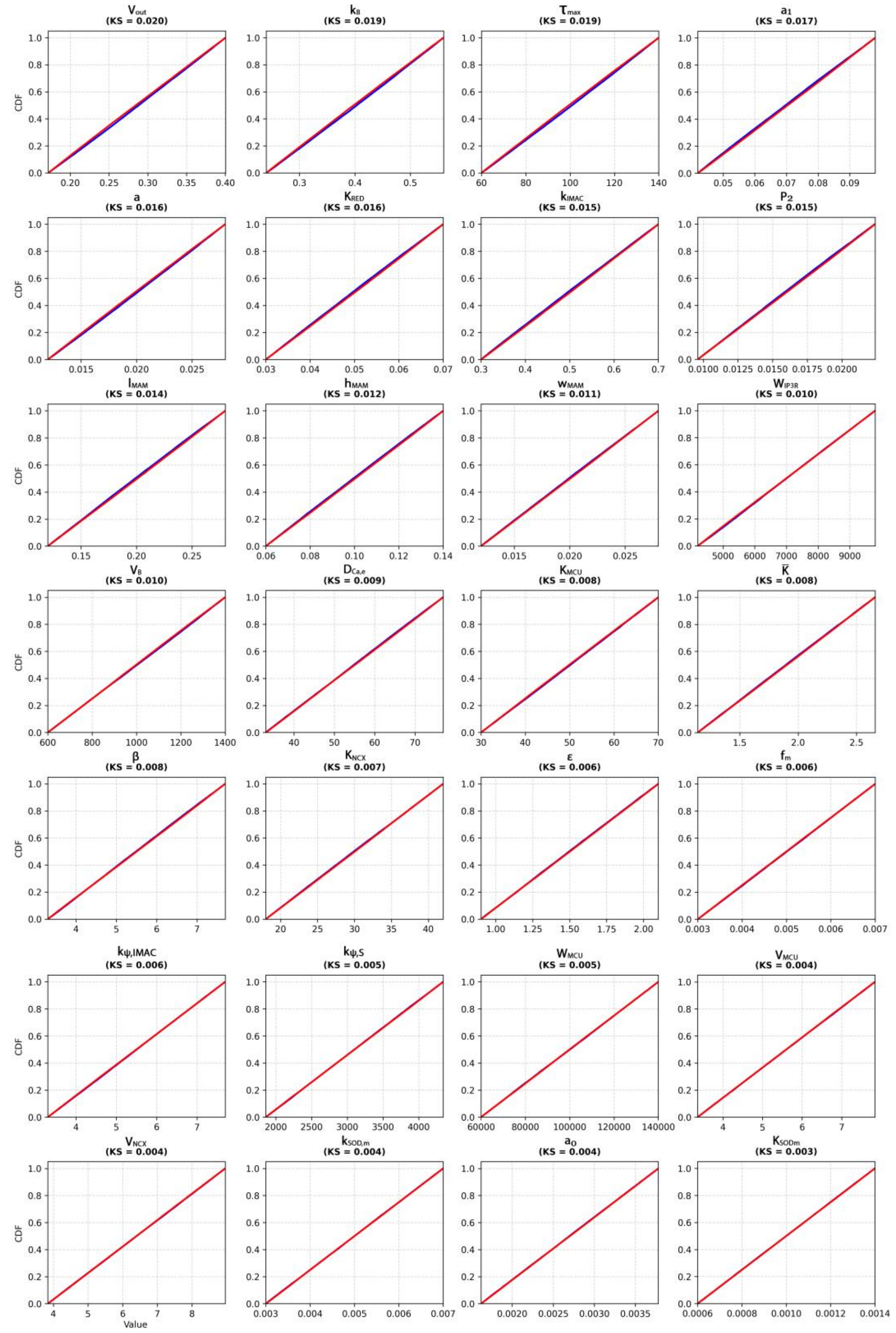


*Fig S11: Continue of Fig S10.*

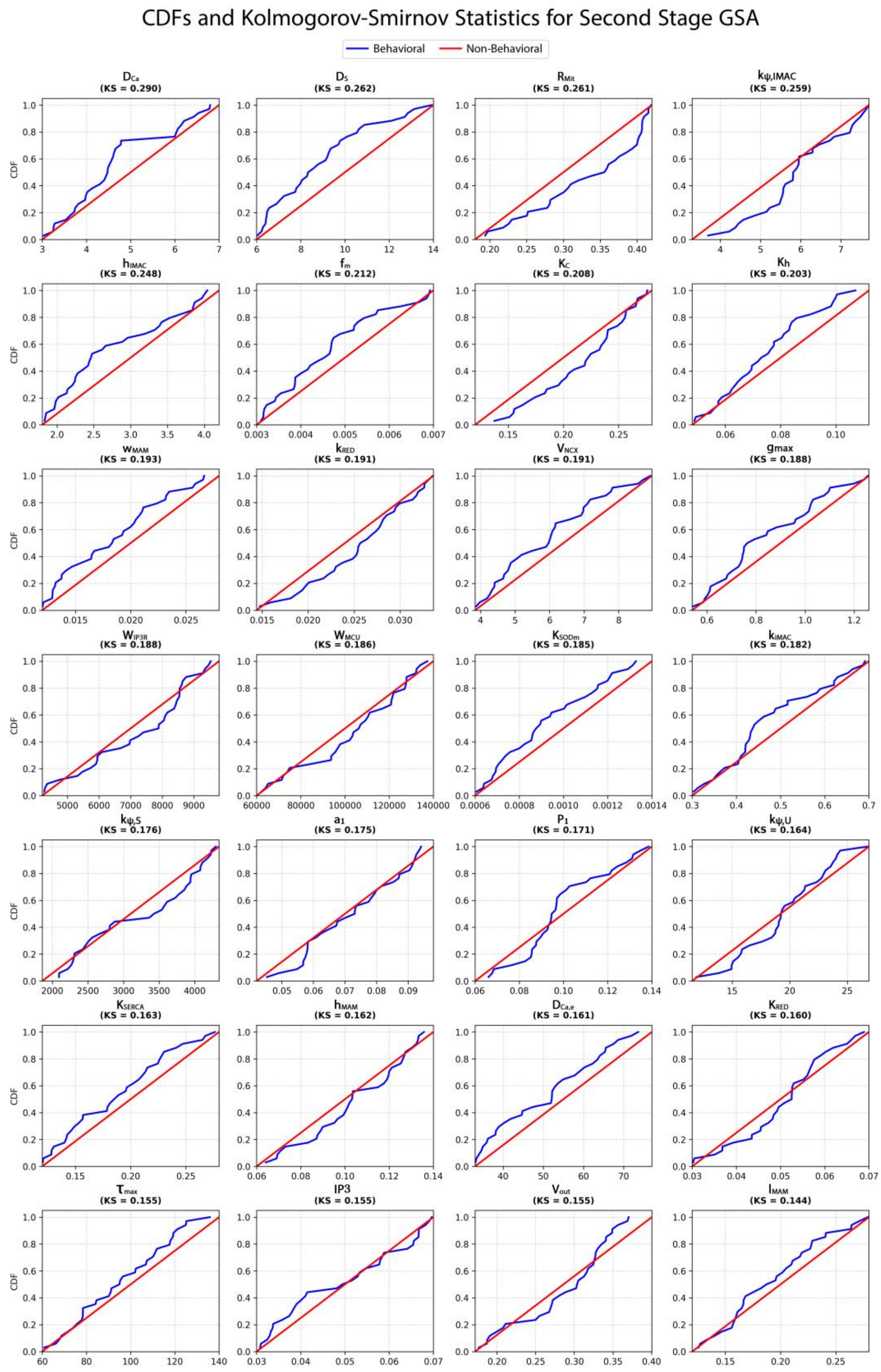


*Fig S12: Cumulative distribution functions plots and Kolmogorov–Smirnov statistics for second-stage global sensitivity analysis of model parameters.*

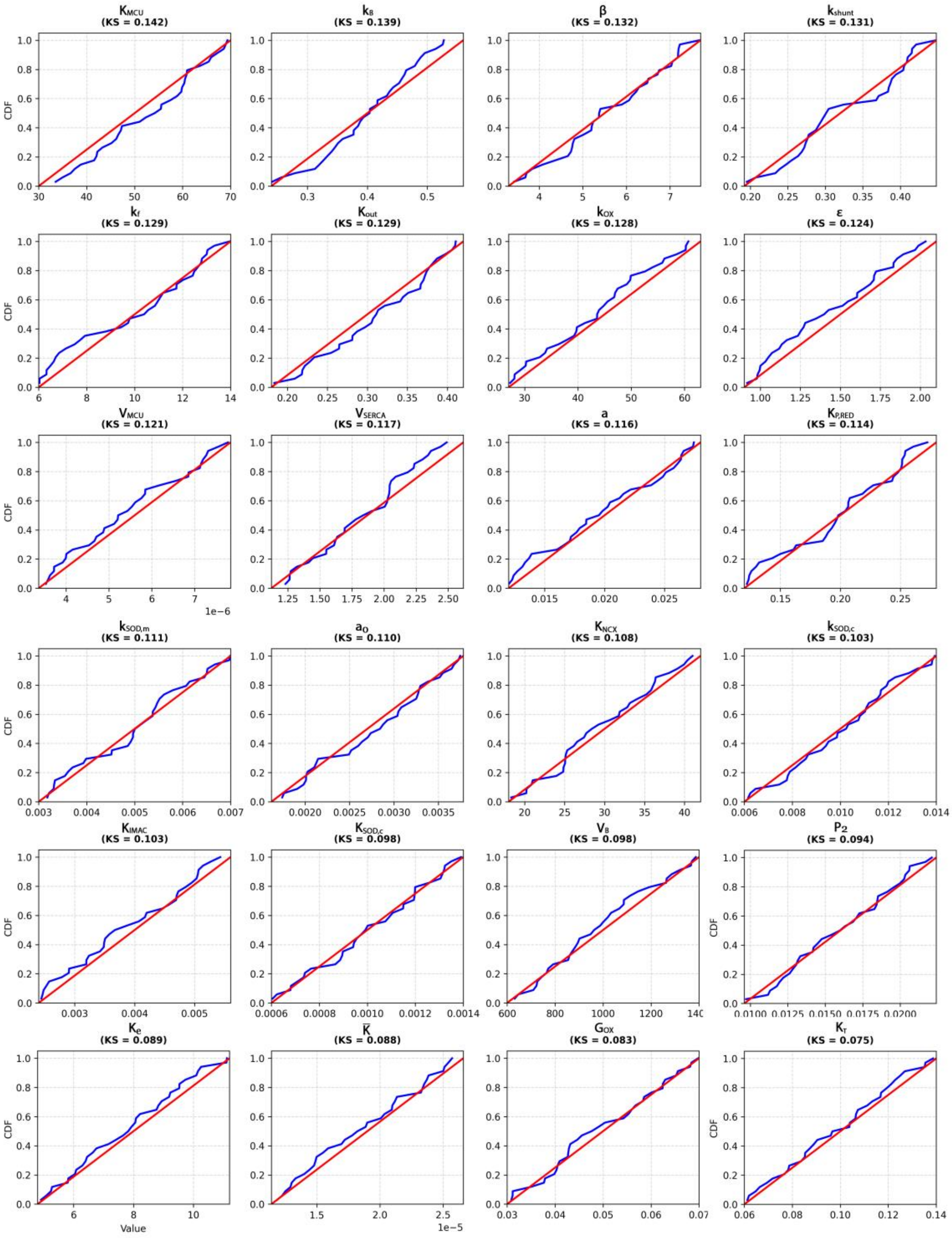


*Fig S13: continue of Fig S12.*

### References


1. Wacquier B, Combettes L, Van Nhieu GT, Dupont G. Interplay Between Intracellular Ca2+ Oscillations and Ca2+-stimulated Mitochondrial Metabolism. Sci Rep. 2016 May 10;6(1):19316.

2. Moshkforoush A, Ashenagar B, Tsoukias NM, Alevriadou BR. Modeling the role of endoplasmic reticulum-mitochondria microdomains in calcium dynamics. Sci Rep. 2019 Dec;9(1):17072.

3. Pages N, Vera-Sigüenza E, Rugis J, Kirk V, Yule DI, Sneyd J. A Model of $$\hbox {Ca}^{2+}$$ Dynamics in an Accurate Reconstruction of Parotid Acinar Cells. Bull Math Biol. 2019 May;81(5):1394–426.

4. Sneyd J, Han JM, Wang L, Chen J, Yang X, Tanimura A, et al. On the dynamical structure of calcium oscillations. Proc Natl Acad Sci. 2017 Feb 14;114(7):1456–61.

5. Watanabe S, Ilieva H, Tamada H, Nomura H, Komine O, Endo F, et al. Mitochondria-associated membrane collapse is a common pathomechanism in SIGMAR1- and SOD1-linked ALS. EMBO Mol Med. 2016 Dec;8(12):1421–37.

6. Wang W, Fang H, Groom L, Cheng A, Zhang W, Liu J, et al. Superoxide Flashes in Single Mitochondria. Cell. 2008 Jul;134(2):279–90.

7. van Stroe-Biezen SAM, Everaerts FM, Janssen LJJ, Tacken RA. Diffusion coefficients of oxygen, hydrogen peroxide and glucose in a hydrogel. Anal Chim Acta. 1993 Feb 15;273(1):553–60.

8. Chalmers S, McCarron JG. The mitochondrial membrane potential and Ca2+ oscillations in smooth muscle. J Cell Sci. 2008 Jan 1;121(Pt 1):75–85.

9. Yang L, Korge P, Weiss JN, Qu Z. Mitochondrial Oscillations and Waves in Cardiac Myocytes: Insights from Computational Models. Biophys J. 2010 Apr 21;98(8):1428–38.

10. Johri A, Chandra A. Connection Lost, MAM: Errors in ER–Mitochondria Connections in Neurodegenerative Diseases. Brain Sci. 2021 Nov;11(11):1437.

11. Wacquier B. Interplay Between Intracellular Ca2+ Oscillations and Ca2+-stimulated Mitochondrial Metabolism. Sci Rep. :16.
